# The adhesion GPCR ADGRV1 controls glutamate homeostasis in hippocampal astrocytes supporting neuron development: first insights into to pathophysiology of *ADGRV1*-associated epilepsy

**DOI:** 10.1101/2024.04.25.591120

**Authors:** Baran E. Güler, Mark Zorin, Joshua Linnert, Kerstin Nagel-Wolfrum, Uwe Wolfrum

## Abstract

ADGRV1 is the largest member of adhesion G protein-coupled receptor (aGPCR) family. In the cell, aGPCRs have dual roles in cell adhesion and signal transduction. Mutations in *ADGRV1* have been linked not only to Usher syndrome (USH), which causes deaf-blindness, but recently also to various forms of epilepsy. While the USH defects are attributed to the loss of fiber links between membranes formed by the extracellular domain of ADGRV1, the pathomechanisms leading to epilepsy remain elusive to date.

Here, we study the specific functions of ADGRV1 in astrocytes where it is highest expressed in the nervous system. Affinity proteomics showed the interaction of ADRGV1 with proteins enriched in astrocytes. Dysregulations of cellular processes important in astrocyte function were indicated by the different transcriptomes of patient-derived cells and Adgrv1-deficent mouse hippocampi compared to appropriate controls. Alteration in morphology and reduced numbers of astrocytes in the hippocampus of Adgrv1-deficent mice. Monitoring the glutamate uptake in colorimetric assay and by live cell imaging of a genetic glutamate reporter consistently showed that glutamate uptake from the extracellular environment is significantly reduced in Adgrv1-deficent astrocytes. Expression analyses of key enzymes of the glutamate-glutamine cycle in astrocytes and the glutamate metabolism indicated imbalanced glutamate homeostasis in Adgrv1-deficient astrocytes. Finally, we provide evidence that the supportive function of astrocytes in neuronal development also relies on ADGRV1 expression in astrocytes. Our data collectively provides first insights into the molecular pathophysiology underlying the development of epilepsy associated with mutations in *ADGRV1*.

**Highlights:**

- ADGRV1 deficiency reduces the number of astrocytes in CA1 and changes the morphology of astrocytes in the hippocampus.
- ADGRV1 interacts with numerous proteins enriched in astrocytes.
- Differential transcriptomes revealed differential expression of genes related to glutamate homeostasis and epilepsy in ADGRV1 deficient models.
- ADGRV1 controls glutamate uptake and regulates homeostasis in astrocytes.
- ADGRV1 in astrocytes is vital for neuron morphogenesis.
- First insights into the molecular pathophysiology underlying the development of epilepsy associated with mutations in ADGRV1.

**Graphical Abstract:** 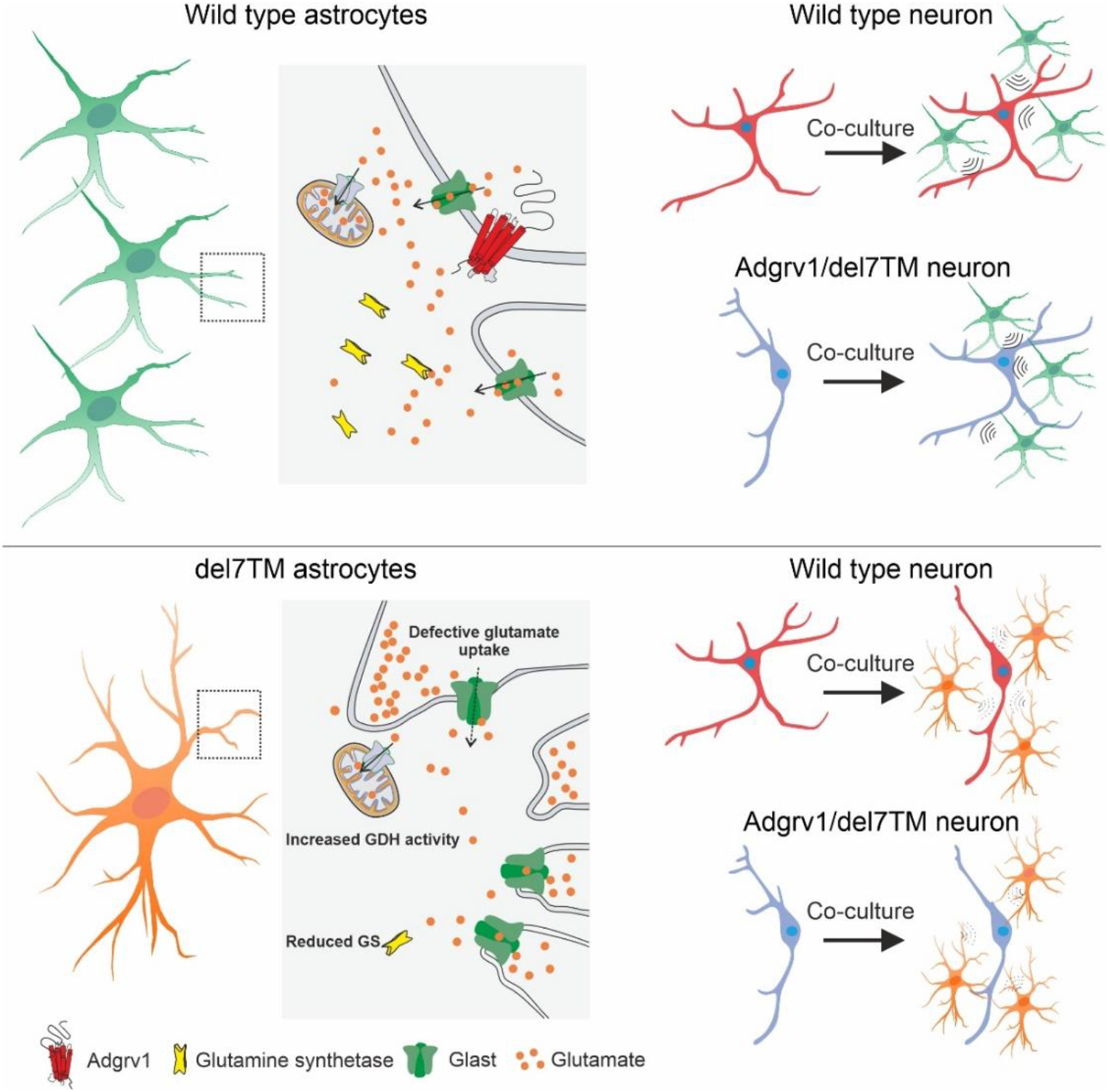

## INTRODUCTION

ADGRV1 (previously known as GPR98, MASS1, or VLGR1) with a molecular weight of approximately 700 kDa is the largest member of the adhesion G protein-coupled receptor (aGPCR) family consisting of 33 other members (Hamann et al., 2015). Adhesion GPCRs are structurally chimeric and contain the signature domains of seven transmembrane receptors and adhesion proteins (Langenhan, 2020). This repertoire makes them suitable to participate in cell adhesion and signaling through G protein coupled pathways (Knapp et al., 2022). Like the other aGPCRs, ADGRV1 can be autoproteolytically processed at its GAIN domain and operate as a metabotropic mechanoreceptor (Kusuluri et al., 2021). Furthermore, there is evidence that mechanosensation of ADGRV1 may trigger its cleavage and subsequent receptor activation by the tethered “Stachel” agonist inducing the switch from Gαs-to Gαi-mediated signaling (Knapp et al., 2022). ADGRV1 is a component of fibrous membrane-links, membrane contacts to the extracellular matrix, namely focal adhesions and in organelle contact sites such as the mitochondria-associated ER membranes (MAMs) (Maerker et al., 2008; Kusuluri et al., 2021; Krzysko et al., 2022; Güler et al., 2023a). There, ADGRV1 were related to the control of cell spreading and migration and the maintenance of Ca^2+^-homeostasis.

In vertebrate hair cells and photoreceptor cells, ADGRV1 forms fiber connections between adjacent membranes, respectively and their loss leads to USH2C, a subtype of the human Usher syndrome, the most common form of hereditary deaf blindness (Weston et al., 2004; Michalski et al., 2007; Maerker et al., 2008). However, in recent years pathogenic variants in *ADGRV1* have been also associated with various forms of epilepsy (Wang et al., 2015; Myers et al., 2018a; Liu et al., 2020; Leng et al., 2022; Zhou et al., 2022). In cohorts of patients with confirmed epilepsy, the rate of *ADGRV1* pathogenic variants was determined to be nearly 3% (Leng et al., 2022). Although most of these epilepsy cases related to *ADGRV1* haploinsufficiency have good prognoses, a recent case report of a heterozygous missense mutation of *ADGRV1* (c.5785G> T) in a patient resulted in hippocampal sclerosis characterized by neuronal cell loss and sudden unexpected death (Ji et al., 2023). It is generally accepted that the epileptic seizures which are characteristic for epilepsy are induced by excessive and abnormal neuronal activity in the brain (Bou Assi et al., 2017). However, the molecular mechanisms leading to the different forms of epilepsy remain elusive to date (Chong et al., 2023; Johannesen et al., 2023). Also, to date, nothing is known about the molecular and cellular pathomechanisms that lead to the epileptic brain conditions caused by defects in ADGRV1.

Although ADGRV1 is expressed almost ubiquitously in many tissues, its expression is strongest in the developing and mature nervous system (McMillan and White, 2010). Of all brain cells, ADGRV1 is most strongly expressed in astrocytes (https://www.proteinatlas.org/). Astrocytes are glia cells and one of the most abundant cells in the central nervous system (CNS). Astrocytes have many roles including, regulation of extracellular ion concentration, control of energy metabolism, glutamate clearance, neuroprotection, synaptogenesis, and establishment of the blood-brain barrier (Mahmoud et al., 2019; Michinaga and Koyama, 2019). The branching of astrocytes promotes extensive contacts with neighboring cells, in particular neurons and specific association with synapses (Higashi et al., 2001). In tripartite synapses, astrocytes wrap around the pre- and post-synapses of neurons and regulate the crosstalk between synapses by balancing the ion and glutamate homeostasis (Perea et al., 2009; Allen and Eroglu, 2017; Broadhead et al., 2022). Glutamate uptake by astrocytes clears the glutamate spillover from synaptic clefts and is part of the glutamate-glutamine cycle between astrocytes and neurons preventing glutamate toxification of the neural tissue (Rose et al., 2018; Mahmoud et al., 2019). However, the uptaken glutamate can be also metabolized in the astrocytes for energy production in the mitochondrial tricarboxylic acid cycle (TCA) (Rose et al., 2020). The dysregulation of both the glutamate-glutamine cycle and the glutamate metabolism in astrocytes can cause an imbalance in brain glutamate homeostasis and lead to hyperexcitability of neurons as seen in epilepsy (Alcoreza et al., 2021). Loss of astrocyte functions or defects in their migratory capacity during development, or postnatal dysfunctions of astrocytes, have been associated not only with neurodegenerative diseases, such as Alzheimeŕs and Parkinson’s disease, but also with the development of various forms of epilepsy (Steinhäuser et al., 2012; Zhang et al., 2016; González-Reyes et al., 2017).

Although ADGRV1 is abundantly expressed in astrocytes, little is known about its specific function in astrocytes of the CNS. In previous studies on ADGRV1 function we made use of primary astrocytes-derived from murine brain as cellular models without a specific focus on the possibly epilepsy-relevant functions in the brain (Kusuluri et al., 2021; Krzysko et al., 2022; Güler et al., 2023b). Our studies collectively revealed insights into a variety of roles of ADGRV1 in cell functions such as cell adhesion, cell migration, mechanosensation, and the regulation of the Ca^2+^-homeostasis at MAMs.

Our current work is focused on elucidating and understanding the specific functions of ADGRV1 in astrocytes by omics, immunohisto- and cytochemistry, biochemistry and a spectrum of cell physiology assays utilizing Adgrv1-deficient mice and human patient cells as models. We show that astrocyte morphology is altered, and their number is significantly reduced especially in the CA1 region of the hippocampus of the Adgrv1-deficient mouse brain. Omics data support the association of ADGRV1 with proteins and processes specifically enriched in astrocytes. Our data also indicates an imbalance in glutamate homeostasis in hippocampal Adgrv1-deficient astrocytes resulting in a reduced glutamate uptake from the extracellular environment and an alteration in the glutamate-glutamine cycle and glutamate metabolism. Our work in primary cultures suggests that ADGRV1 is also important for the proper development of neurons and showed that ADGRV1 substantially contributes to the beneficial support of astrocytes for neurons. Overall, our data provides first insights in the molecular pathophysiology underlying the development of epilepsy associated with mutations in *ADGRV1*.

## Material and Methods

### Animals

Experiments were conducted in accordance with the guidelines set forth by the Association for Research in Vision and Ophthalmology. Mice were housed in a controlled environment with a 12/12-hour light/dark cycle with unrestricted access to food and water. Adgrv1/del7TM mice, susceptible to audiogenic seizures, carry a nonsense mutation, V2250X, in the *Adgrv1* gene, leading to the loss of both its transmembrane and cytoplasmic domains (McMillan and White, 2004). The mice were bred on a C57BL/6 background. The use of mice for research purposes was granted approval by the District Administration Mainz-Bingen under reference number 41a/177-5865-§11 ZVTE on April 30, 2014.

### Antibodies

The following antibodies were used: rabbit anti-GFAP (DAKO agilent, ZO334), rabbit anti-SOX9 (Abcam, ab185966), guinea pig anti-MAP2 (Sysy, 188004), rabbit anti-GLAST (Almone lab, AGC-021), mouse anti-Gapdh (Abcam, ab9484), rabbit anti-Glutamine synthetase (Abcam, ab49873), mouse anti-Glutamine synthetase (Abcam, ab64613), rabbit anti-Homer1 (Sysy, 160011), mouse anti-Gephyrin (Sysy, 147021). Secondary antibodies conjugated to Alexa488, Alexa568 and Alexa647 were purchased from Molecular Probes (Life Technologies). Nuclear DNA was stained with DAPI (4′,6-diamidino-2-phenylindole, 1 mg/ml, Sigma-Aldrich, 10236276001).

### Transcardiac perfusion fixation of mouse brains and immunohistochemistry analysis

Mice were anesthetized with an injection of 40 to 80 mg/kg of pentobarbital. Then transcardiac perfusion was performed with cold 0.1 M phosphate buffer (pH 7.4) (PBS) containing 0.0025 g/50 ml heparin solution following by 4% paraformaldehyde (PFA) in phosphate buffer (Gage et al., 2012). Dissected brains were post-fixed in buffered 4% PFA overnight, infiltrated stepwise with 10%, 20% and, 30% sucrose in PBS, and embedded in Tissue-Tek OCT before frozen in melting isopentane (Wolfrum, 1991). 16 µm thick cryosections and placed onto Poly-L-Lysine (Sigma-Aldrich, P4832-50ML) coated coverslips. After permeabilization in 0.2% Triton X-100 for 10 min (Sigma-Aldrich, 102533092) in PBS, immunostaining performed as previously described (Arévalo et al., 2022). Fluorescence intensity immunoreactivity in mouse hippocampi was analyzed using a build-in ImageJ/Fiji plugin (https://fiji.sc/).

### Morphometric analysis of hippocampal astrocytes

The morphology of astrocytes and neurons was quantified as previously published (Young and Morrison, 2018). In brief, cryosections through the hippocampus of mice were stained for astrocytes by anti-GFAP and counterstained with DAPI. Hippocampal subregions were identified by the Allen mouse brain atlas (https://mouse.brain-map.org/) (Figure S1). Z-stacked images were maximum intensity projected, converted to 8-bit using ImageJ/Fiji, and pre-processed with an FFT bandpass filter (filter up to 5 pixels, down to 40). Individual astrocytes were cropped, skeletonized, and quantified using AnalyzeSkeleton(2D/3D) and FracLac analysis plugins (Arganda-Carreras et al., 2010).

### Astrocyte number analysis

GFAP and SOX9 positive astrocyte numbers were counted in different hippocampal subregions in immunostained cryosections of mice brains. GFAP and DAPI positive cells were marked with the image counter plugin of ImageJ/Fiji. For the analysis of SOX9-positive cells, images were converted to 8-bit and “huang dark” threshold was applied, converted to binary, and watershed was applied. Particles analysis was performed with the sizes defined as 100 to infinity and circularity was defined as 0.4 to 1.00. Counted cells were exported and processed in GraphPad prism.

### Mouse hippocampi transcriptome analysis

PN40 wild type and Adgrv1/del7TM mice were sacrificed by cervical dislocation. The hippocampus was dissected from the brain and flash-frozen using liquid nitrogen. mRNA was isolated according to the instructions of the Qiagen RNeasy Mini kit. The quality and quantity of RNAs were measured with a Nanodrop 2000 spectrophotometer (Thermo Fischer Scientific). Sequencing was performed with Illumina NovaSeq 6000 Sequencing Systems (Novogene). Analysis was performed using Conda package manager v.23.5.2 (Grüning et al., 2018). All packages were installed in separate environments and an analysis pipeline was written in Snakemake v.7.32.3 (Mölder et al., 2021). First, we performed a quality control analysis of raw paired end reads using FastQC v.0.12.1 (https://www.bioinformatics.babraham.ac.uk/projects/fastqc/). Then we trimmed our reads using Trimmomatic v.0.39 (Bolger et al., 2014) with HEADCROP:15 parameter. After trimming, we performed contamination check analysis using FastQ Screen v.0.15.3 (Wingett and Andrews, 2018) software and reference genomes: *Escherichia coli* strain K-12 (ASM584v2), *Homo sapiens* (GRCh38.p14), *Metamycoplasma orale* strain NCTC10112 (50465_D02-3), *Mus musculus* (GRCm39), *Staphylococcus aureus* (ASM1342v1). During contamination check, we used Bowtie2 v.2.5.1 as an aligner. Reads that did not match to any mammalian genome or aligned to *E. coli*, *M. orale* or *S. aureus* were excluded from subsequent analysis. Then, we aligned the filtered reads to the reference mouse genome (GRCm39) using Hisat2 v.2.2.1 (Kim et al., 2019) software, sorted and recorded them in the bam format using Samtools v.1.17 (Li et al., 2009). To quantify the number of reads that mapped to each gene we used the featureCounts program that included in Subread v.2.0.6 (Liao et al., 2014) package. In the following analysis we used only genes with more than 5 mapped reads. After filtering files, we performed a differential expression analysis using DESeq2 v.1.40.2 (Love et al., 2014). In further analysis we used only genes with adjusted p-value less than 0.05. Three biological replicates of WT and Adgrv1/del7TM mice were used. However, during the analysis of the raw data, we observed high ingroup variation in Adgrv1/del7TM. Therefore, to identify the differences between WT and Adgrv1/del7TM group, we excluded the outlier biological replicate to decrease ingroup variation.

### Human fibroblast transcriptome analysis

Dermal primary fibroblast lines were expanded from skin biopsies of human subjects (ethics vote: Landesärztekammer Rhineland-Palatinate to KNW). Fibroblasts from the skin biopsies of a 57-year-old male patient with clinically confirmed Usher syndrome 2C caused by a nonsense mutation in the *ADGRV1* gene (g.[90006848C>T], R2959*) were kindly provided by Dr Erwin van Wijk (Radboud University Medical Center, Nijmegen). Fibroblasts derived from the patient and from a healthy individual were maintained in DMEM/10% FBS (Thermo Fischer Scientific, 31966-021) before the RNA isolation, isolated fibroblast RNAs were subjected to RNAseq analysis with Illumina NovaSeq 6000 Sequencing Systems (Novogene) previously described (Schäfer et al., 2023).

### Analysis of tandem affinity purification (TAP) data sets and Gene Ontology (GO) analysis

Data sets obtained by tandem affinity purification (TAP) from HEK293T cells lysates expressing Strep II/ Strep-Tactin tagged ADGRV1 constructs (Figure S2A, B) were previously published in (Knapp et al., 2022). We categorized the TAP hits by using Gene Ontology (GO) term analysis following by Cytoscape (http://www.cytoscape.org; accessed on 09 September 2023) plugin ClueGO (Bindea et al., 2009). Functional protein association networks were prepared by using String database v10 (Szklarczyk et al., 2015). For transmembrane transporter activity related genes in fibroblasts were screened in the web application of the Amigo2 (https://amigo.geneontology.org/amigo/search/bioentity?q=transmembrane%20transporter%20activity).

### Isolation of primary astrocytes

Astrocytes were isolated from newborn female and male sibling mouse pups on postnatal day 0 (PN0), as previously described (Güler et al., 2021). Briefly, PN0 mouse pups were dissected, and their brains enzymatically and mechanically dissociated. The resulting single-cell mixtures were plated on Poly-L-Lysine-coated (PLL) flasks and cultured in DMEM/10% FBS/2% penicillin/streptomycin for 7 to 10 days. Upon reaching confluence, flasks were agitated to remove oligodendrocytes and neurons. To separate primary astrocytes from microglia cells, cultures were trypsinized and plated on sequential bacterial-grade dishes. Microglia cells adhered to the dish surfaces, while astrocytes were recovered from the supernatant. Isolated primary astrocytes were cultured for an additional 7 to 10 days in complete growth medium.

### Isolation of primary neurons

Primary neurons were isolated from the mouse hippocampus on embryonic day E18.5. Hippocampal tissues were dissected and kept in ice-cold HBSS buffer. After washing twice with HBSS to remove dead cells, tissues were trypsinized for 5 min at 37°C. Single-cell solutions were prepared using fire-polished Pasteur pipettes with varying opening diameters. After trituration, 5 ml HBSS was added to stop the enzymatic reaction. Cell suspensions were allowed to settle for 5 min before aspirating the HBSS/trypsin mixture. Cell pellets were then triturated with 2 ml cold HBSS, and cell numbers were counted. Cells were seeded on PLL coated coverslips and cultured in neuron growth medium containing B27 supplement, L-glutamine, penicillin-streptomycin, and glutamate in Neurobasal medium (ThermoFisher Scientific, 21103049).

### Astrocyte-neuron co-culture

Primary astrocytes were cultured in PLL coated coverslips in a 6 well plate for 2 days at a density of 3x10^5^ cells per well in DMEM/10% FBS/2% penicillin/streptomycin medium. After two days of cultivation, primary neurons were isolated and seeded on top of the lawn of astrocytes at a density of 3x10^5^ neurons per well. In these co-cultures, the DMEM medium was replaced by neuron growth medium.

### Immunocytochemistry

Primary astrocytes and neurons were processed for immunocytochemistry as previously described (Kusuluri et al., 2021; Güler et al., 2023b).

### Western blot analyses

Protein lysates from tissue and primary cells were prepared as previously described (Krzysko et al., 2022) and densitometry analysis was performed by using an Odyssey imaging system (LI-COR Biosciences).

### Glutamate uptake and Glutamate dehydrogenase (GDH) activity assays

Glutamate uptake assays were performed as previously described (Mahmoud et al., 2019). Briefly, primary astrocytes were seeded at a density of 2x10^5^ cells per well of 12 well plate. After 72 h in DMEM medium, astrocytes were treated with serum free Opti-MEM medium for 2 h. Opti-MEM was removed by 3 PBS washes. Cells were exposed to 100 or 200 µM Glutamate solution in Hank’s Balanced Salt Solution (HBSS) containing Ca^2+^ buffer for 4 h. Cell culture supernatants were analyzed in a Glutamate assay kit (Sigma-Aldrich, MAK004) following the manufacturer’s instructions. GDH activities in primary astrocytes were determined with glutamate dehydrogenase activity assay kit following with the manufactureŕs instructions (Abcam, ab102527).

### Stachel peptide synthesis

The “Stachel” peptide (SVYAVYARTDN) (Knapp et al., 2022) and a randomized control peptide (ATSVRYDNAYV) were synthesized from C-to N-terminus by solid phase peptide chemistry, HPLC purified and analyzed by mass spectrometry (Chempeptide limited). Lyophilized peptide was solubilized in 100% DMSO and diluted in 80% ddH_2_0 and 20% HNO_3_ (20% v/v) to obtain a 10 mM stock solution.

### Live cell imaging and “Stachel” peptide application

Glutamate was monitored in primary astrocytes using the fluorescent reporter for glutamate pAAV.GFAP.iGluSnFR3.v857.GPI (Addgene plasmid # 178338) which was a gift from Kaspar Podgorski (Aggarwal et al., 2023). Images were acquired by Nikon eclipse Ti2 microscope (Nikon Instruments Inc) equipped with CSU-W1-T2 automatic Spinning Disk Confocal Scanning unit with 63x water objective (Yokogawa Electric Corporation). Dyes were excited with a 488 nm laser. The exposure time was kept at 250 ms and image sequences were obtained every 700 milliseconds for a total of 300 seconds. 1 mM “Stachel” peptide applied to the HBSS medium to evoke the Adgrv1 signaling at 98^th^ sec of imaging. Fluorescence values were normalized to the first 98 frames of images (F_0_). Calculation of intensity changes was done by following formula (F-F0)/ F_0_.

### Morphometric analysis of neurons

To analyze neuronal dendritic complexity, Sholl analysis method was used as previously described (Ferreira et al., 2014). Briefly, images of MAP2 positive neurons were exported to ImageJ/Fiji and converted to 8-bit images and “IsoData” thresholding was applied to images. Thresholded images were converted to Mask and fill hole function applied. Binarized images were loaded into the SNT plugin in ImageJ/Fiji. Branches of neuritis were selected and semi-autonomous branch detection option of SNT used. Intersection numbers were exported to Excel and Sholl plots were generated using R Studio.

### Synaptic puncta analysis in cultured neurons

Primary neurons isolated from WT and Adgrv1/del7TM mice hippocampi were cultured for 14 days. For the quantification of the number of synaptic puncta along the MAP2 positive dendrites, SynapCountJ and NeuronJ Fiji/ImageJ plugins were used (Mata et al., 2016). Briefly, dendrites from individual neurons were tracked using NeuronJ plugin and dendritic structures saved as a tab-limited text file (Txt). Next, dendritic branch information and images with synaptic channels were uploaded to SynapCountJ plugin. Images were manually thresholded and threshold values kept constant throughout the analysis. Synaptic densities were calculated per 100 µm dendritic length. Synaptic puncta sizes were analyzed with another Fiji/ImageJ plugin Synapse Counter (Dzyubenko et al., 2016). Images were thresholded with Triangle and synaptic particle size was limited from 20 px^2^ to 400 px^2^ using the batch mode function of the plugin.

### Microscopy

Images were acquired using a Leica DM6000B Microscope (Leica-Bensheim) or a Nikon eclipse Ti2-E microscope (Nikon Instruments Inc) equipped with CSU-W1-T2 automatic Spinning Disk Confocal Scanning unit (Yokogawa Electric Corporation).

### Statistical analyses

Statistical analyses were performed with Graphpad Prism 9.0 software (GraphPad Software Inc.). Differences between two sets of data were assessed via a two-tailed Student’s t-test. In the case of multiple group comparisons, one way ANOVA was employed, followed by either Dunnett’s multiple comparison tests or Sidak’s multiple comparison tests, chosen based on the specific data being compared. Significance levels were defined as *p < 0.05, **p < 0.01, and ***p < 0.001. The bar plots display means with standard deviations (mean ± SD). The box plots exhibit the median (central line), while the edges of the boxes represent the interquartile range (25^th^-75^th^ percentile). The whiskers depict the range for the upper 25% and lower 25% of the data points.

## RESULTS

### Adgrv1 controls astrocyte morphology in the mature mouse hippocampus

We have previously shown that the morphology of primary astrocytes is altered in Adgrv1/del7TM mice (Kusuluri et al., 2021). To examine the astrocyte morphology *in situ,* we stained cryosections of the hippocampus of mature Adgrv1/del7TM and control wild-type (WT) mice for the astrocyte marker glial fibrillary protein (GFAP) (Figures 1B, and S1). We then quantified the branching, the circularity, and the convex-hull area of astrocytes in the different subregions of the hippocampus (Figure 1C). The number of branches in astrocytes was significantly increased in all three *Cornu ammonis* regions of the hippocampus (CA1, CA2, and CA3) in Adgrv1/del7TM compared to WT (Figures 1D-F). The circularity (polarization) and convex-hull area of the astrocytes were significantly reduced in CA1 and CA2, but not in CA3 (Figures 1E, F). These data suggest that the lack of Adgrv1 results in morphological changes in astrocytes, predominantly in the CA1 region of the hippocampus.

**Figure 1.**
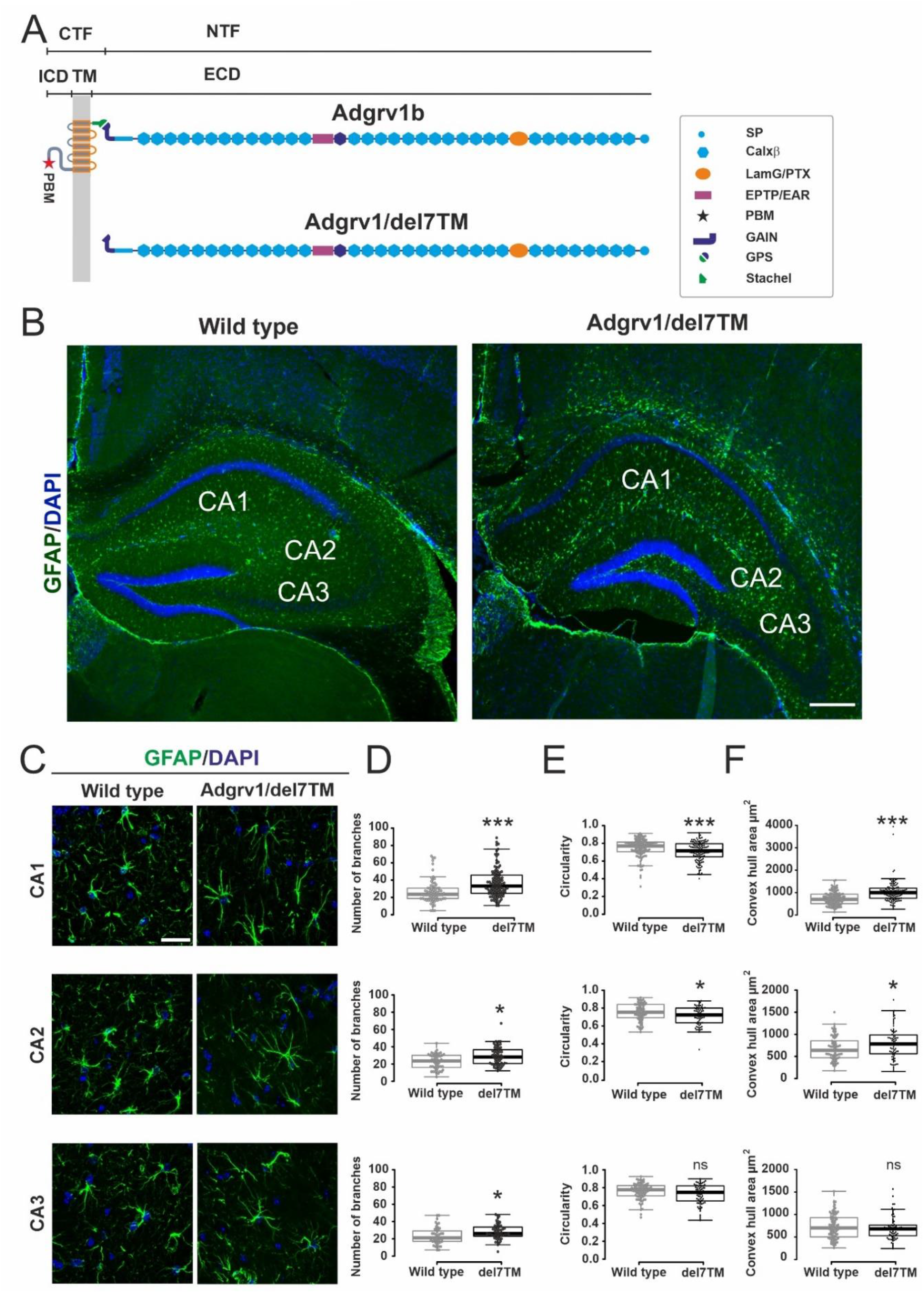
Morphological analysis reveals differences between astrocytes in WT and Adgrv1/del7TM mouse hippocampi. **(A)** Domain structures of ADGRV1 full length and Adgrv1/del7TM proteins. The cleavage of the full-length molecule from the highly conserved GPCR proteolytic site (GPS) in the GAIN (autoproteolysis-inducing) domain results in relative short C-terminal fragment (CTF) which contains 7-transmembrane (7TM) and intracellular domain (ICD) and extra-large N-terminal fragment (NTF). Extra-large N-terminal fragment contains calcium binding Calxß repeats, seven epilepsy-associated/Epimptin like (EAR/EPTP) repeats, and pentaxin/laminin G-like domain (Lamg/PTX). Adgrv1/del7TM mice carry a nonsense mutation in V2260* which results in STOP codon and leads to deletion 7TM and ICD domains. **(B)** Overview of hippocampal sections from WT and Adgrv1/del7TM hippocampus through the CA (*Cornu ammonis*) 1, CA2 and CA3 subregions of the hippocampus stained for GFAP and for nuclear DNA by DAPI. **(C)** Anti-GFAP stained astrocytes in the three different CA1-3 regions of the hippocampus and the quantification of **(D)** branching, **(E)** circularity and **(F)** convex hull areas. n = 3 per mouse group, 3 continuous sections were analysed. In total 150-160 cells from CA1, 80-85 cells from CA2 and 75-85 cells from CA3 were used for the quantification. Scale bar: 25 μm. Statistics: two-tailed Student’s t test; \**p*≤0.05, \*\**p*≤0.01, \*\*\**p*≤0.001.

### Loss of astrocytes in hippocampus regions of Adgrv1/del7TM mice

Next, we examined whether the number of astrocytes in the hippocampus is altered in Adgrv1/del7TM mice. The number estimated by determining the level of expression of the astrocyte-specific markers GFAP and SOX9 in Western blots of hippocampus tissue (Figures 2A, B). Quantification of Western blot bands showed a slight decrease of GFAP and SOX9 expression in the hippocampus of Adgrv1/del7TM mice. We subsequently stained cryosections through the hippocampus for GFAP, SOX9, and DAPI and counted the immunostained astrocytes in the different regions (Figures 2C, D). There were no significant changes in the number of astrocytes in the CA2, CA3, and dentate gyrus region, but a significant decrease in the number of astrocytes in the CA1 region of Adgrv1/del7TM hippocampi compared to WT controls.

**Figure 2.**
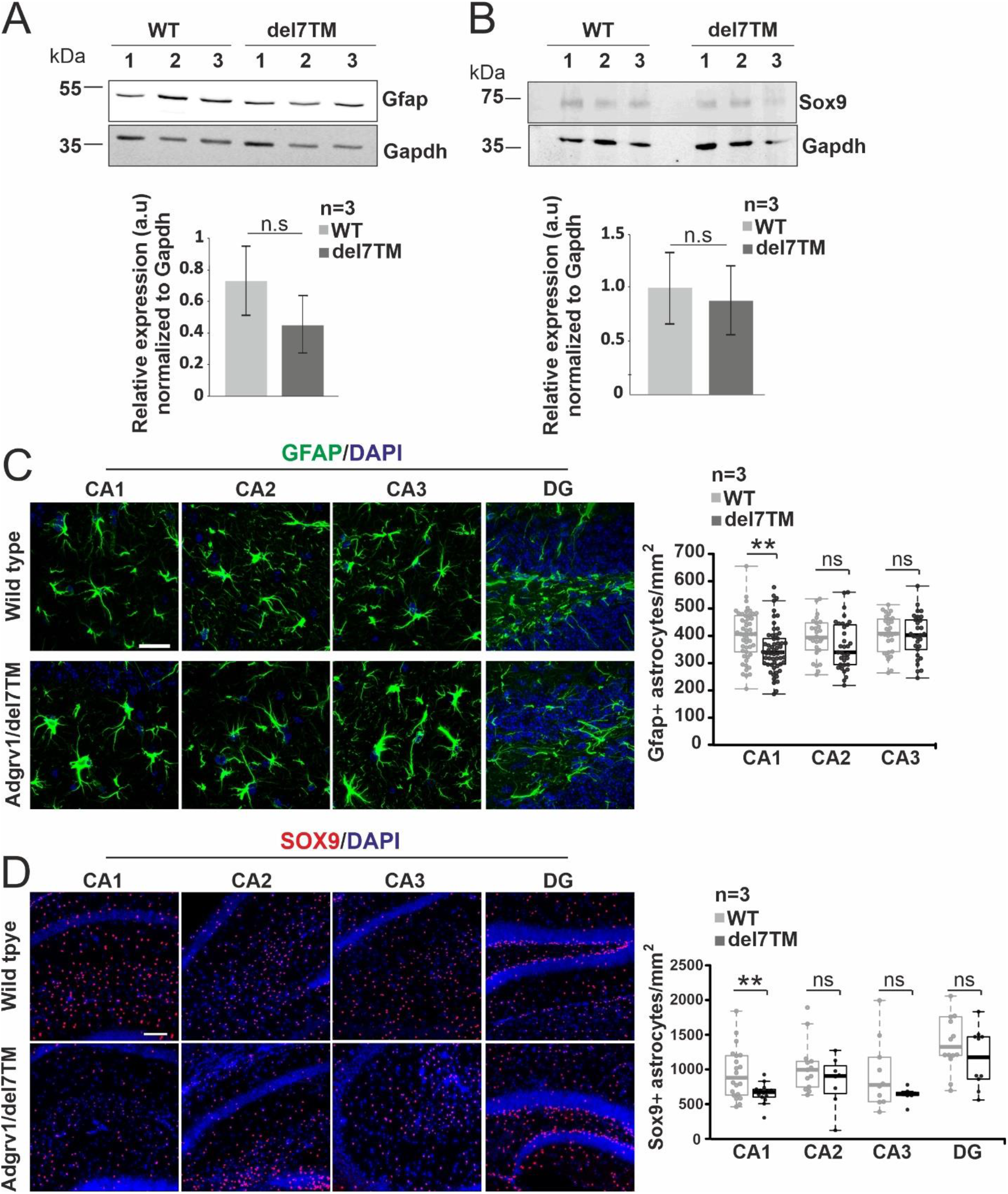
Analysis of the abundance of astrocytes in the hippocampus of Adgrv1/del7TM and wild type mice. **(A, B)** Quantitative Western blot analyses of the astrocyte marker proteins GFAP **(A)** and SOX9 **(B)** demonstrate the decrease of the expression of both proteins in the hippocampus of Adgrv1/del7TM mice. **(C, D)** Immunofluorescence staining of GFAP **(C)** and SOX9 **(D)**, counterstained by DAPI for nuclear DNA in cryosections through the hippocampus of Adrgv1/del7TM and WT control mice. Quantification of GFAP-positive and SOX9-positive astrocytes revealed the significant decrease in the number of astrocytes only in the hippocampus CA1 region of Adgrv1/del7TM mouse compared to WT. n = 3 per mice group, 3 continuous sections were analysed. Scale bars: C: 25 μm; D: 200 µm. Statistical evaluation by two-tailed Student’s t test; \**p*≤0.05, \*\**p*≤0.01, \*\*\**p*≤0.001.

### Affinity proteomics reveals the interaction of ADGRV1 with proteins enriched in astrocytes

We have previously identified numerous putative interacting proteins of ADGRV1 by affinity proteomics capture approaches based on tandem affinity purifications (TAP) combined with mass spectrometry (Figure S2A) (Knapp et al., 2019, 2022). Re-analyzing our TAP datasets previously obtained in HEK293T and hTERT-RPE1 cells (PXD042629), we identified 289 proteins which are enriched in the proteome of astrocytes (Batiuk et al., 2020) as potential interacting proteins to ADGRV1 (Table S1, subtitle 1: Enriched protein annotations; Figure S2B). GO term analysis of these astrocyte proteins showed that in total 52 proteins are clustered in the GO term “*Transmembrane transporter activity*” (Table S1, subtitle 2: Biological function analysis; Figure S2C). Interestingly, five of these potentially interacting proteins of ADGRV1 were also found under the GO term *“L-glutamate import”* (subtitle Table S1; Figure S2C, D) suggesting a link to the glutamate homeostasis, a major function of astrocytes in the CNS. In particular, the two plasma membrane proteins, namely the excitatory amino acid transporters SLC1A1/EAAT3 and SLC1A3/EAAT1/GLAST1 are core proteins of the glutamate uptake machinery in astrocytes (Cuellar-Santoyo et al., 2023) and the three mitochondrial carrier family proteins, SLC25A12/Aralar, SLC25A13, and SLC25A22 are related to the glutamate metabolization in the TCA (Table S1, subtitle 3: Transporters and L-Glutamate import; Figure S2D) (Hillen and Heine, 2020). These findings indicated that ADGRV1 potentially interacts with numerous proteins that are important for the astrocyte functions in the nervous system.

### Transcriptome analysis reveals differentially expressed genes (DEGs) related to glutamate homeostasis and epilepsy in the hippocampus of Adgrv1/del7TM mice

Next, we performed genome-wide mRNA sequencing from hippocampus tissue dissected from Adgrv1/del7TM and WT mice. The comparison of transcriptomes revealed 80 differentially expressed genes (DEGs) in the hippocampus of Adgrv1/del7TM in relation to WT mice (Table S2). The removal of the non-coding RNAs from the list of DEGs due to incomplete genome annotation (Boivin et al., 2020) resulted in 25 and 31 protein-coding genes being significantly up or down regulated, respectively (Figure 3A, B; Table S2).

**Figure 3.**
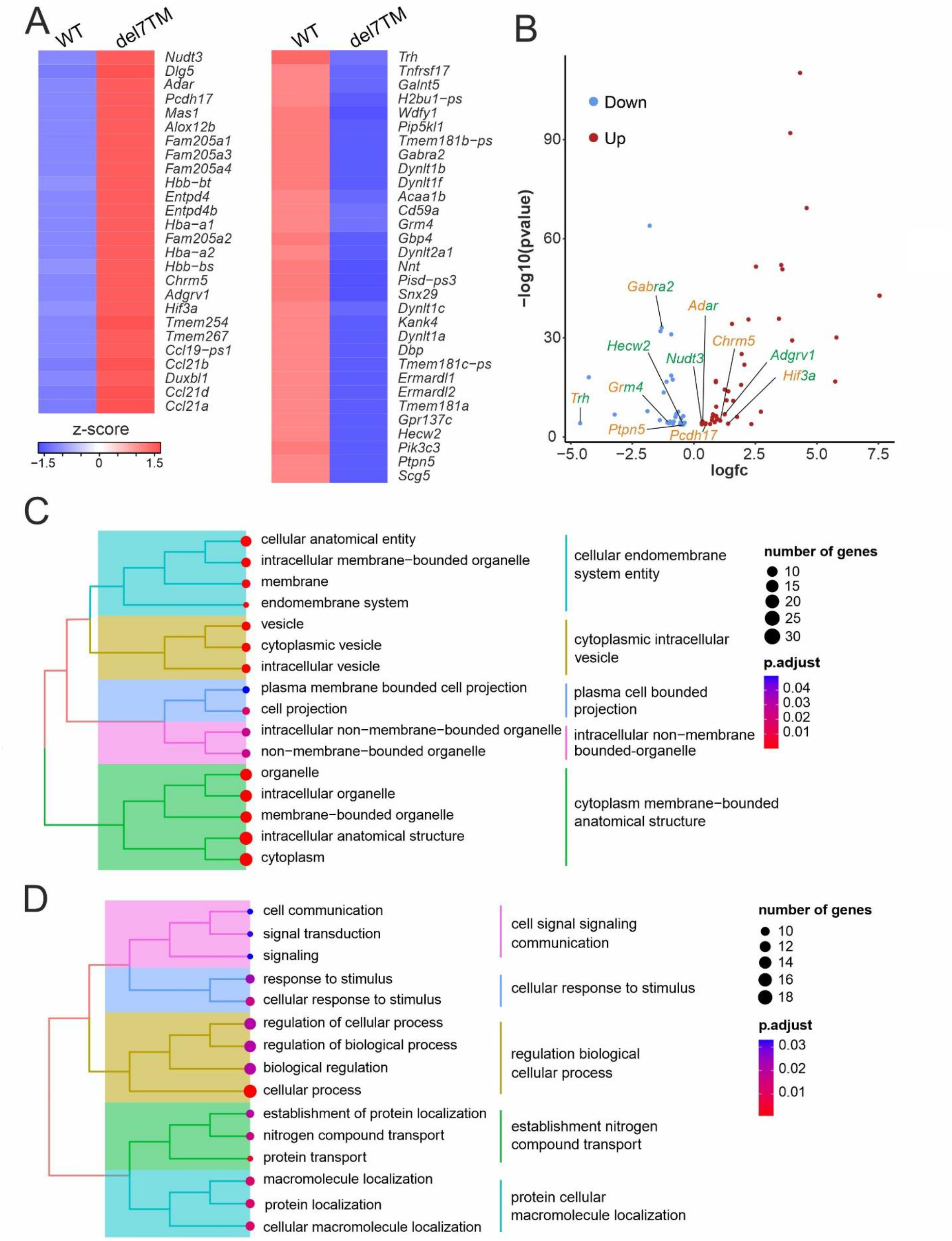
Transcriptome analysis of the hippocampus of Adgrv1/del7TM and WT mice. **(A)** Heatmap of differentially expressed genes (DEG) in the hippocampus of wild type control (WT) and Adgrv1/del7TM (del7TM) mice. Red, upregulated, and blue, downregulated genes in Adgrv1/del7TM compared to WT. Only adjusted p-values less than 0.05 are shown. **(B)** Volcano plot of DEGs associated to epilepsy (green) and related to transporter activity (orange) in the hippocampus Adgrv1/del7TM mice. **(C)** Gene set enrichment (GSE) analysis for the biological function of the DEGs shown in clusters. **(D)** GSE for the cellular compartment analysis of DEGs. *n*=3 biological replicates for WT and *n*=2 biological replicates for del7TM.

Gene set enrichment analysis (GSEA) of the term “*cellular compartment”* showed that DEGs in the hippocampus of Adgrv1/del7TM mice are associated with “*cellular endomembrane system entity*”, “*cytoplasmic intracellular vesicle*”, “*plasma cell bounded projection*”, “*intracellular non-membrane-bounded organelle”* and “*cytoplasm membrane-bounded anatomical structure*” (Figure 3C) (date of analysis: 24.08.2023). GSEA of “*biological function”* showed that DEGs in Adgrv1/del7TM hippocampus are associated with functions related to “*cell signaling*”, “*cellular response to stimulus*”, “*regulation of biological processes*”, “*establishment of nitrogen compound transport*” and “*protein cellular macromolecule localization*” (Figure 3D) (date of analysis: 24.08.2023).

In screens of the Harmonizome 3.0 database (Rouillard et al., 2016) (date of analysis: 22.12.2023) we found 6 DEGs associated with epilepsy diseases in Adgrv1/del7TM hippocampus. Of these genes, 4 were found to be upregulated (*Nudt3*, *Adar*, *Adgrv1*, *Hif3a*) while 2 were downregulated (*Trh*, *Grm4*). Interestingly we found that epilepsy associated genes *Adar, Hif3a, Trh,* and *Grm4* also play a role in glutamate receptor and metabolism regulation. In addition to those DEGs, we identified *Pcdh17, Chrm5, Gabra2,* and *Ptpn5* which function as glutamate receptor and metabolism regulatory proteins (Figure 3B).

### Transcriptome analysis of human patient-derived cells confirmed DEGs related to glutamate homeostasis and epilepsy

Next, we performed genome-wide mRNA sequencing of dermal fibroblasts of a confirmed USH2C patient and a healthy individual by Illumina platform and pair end reads were mapped and quantified (Figure 4). Our DEG analysis of the transcriptomes revealed a total of 1,319 DEGs (Table S3, subtitle 1: All DEGs of patient vs healthy fibroblasts). By GO term analysis applying the web application of the amigo2 and previously reported epilepsy-associated genes (Macnee et al., 2023) we identified in 87 genes related to “transporter activity” and 63 genes related to “epilepsy associated”, respectively (Table S4, subtitle 2: Transporter activity genes, subtitle 3: Epilepsy associated genes). (https://amigo.geneontology.org/amigo) (Figure 4A, B). 24 of these DEGs (15 up-regulated and 9 down-regulated) were identified as being common to genes involved in transporter activity and to genes associated with epilepsy. (Figure 4B, Table S3, subtitle 4: Common in Transporter activity and epilepsy associated gene). For further GSEA analysis, we combined “transporter activity” and “epilepsy associated genes” which was a total number of 125 DEGs. Of these DEGs, 74 genes were up-and 51 genes were downregulated (Figure 4A, S3, Table S3, subtitle 5: DEGs in epilepsy and transport). A comparison with of these DEGs with those found in the Adgrv1/del7TM hippocampus revealed that the solute carrier proteins *SLC2A1, SLC4A3,* and *SLC19A3* were present as DEGs in both transcriptomes (Figure 4A, B, S3).

**Figure 4.**
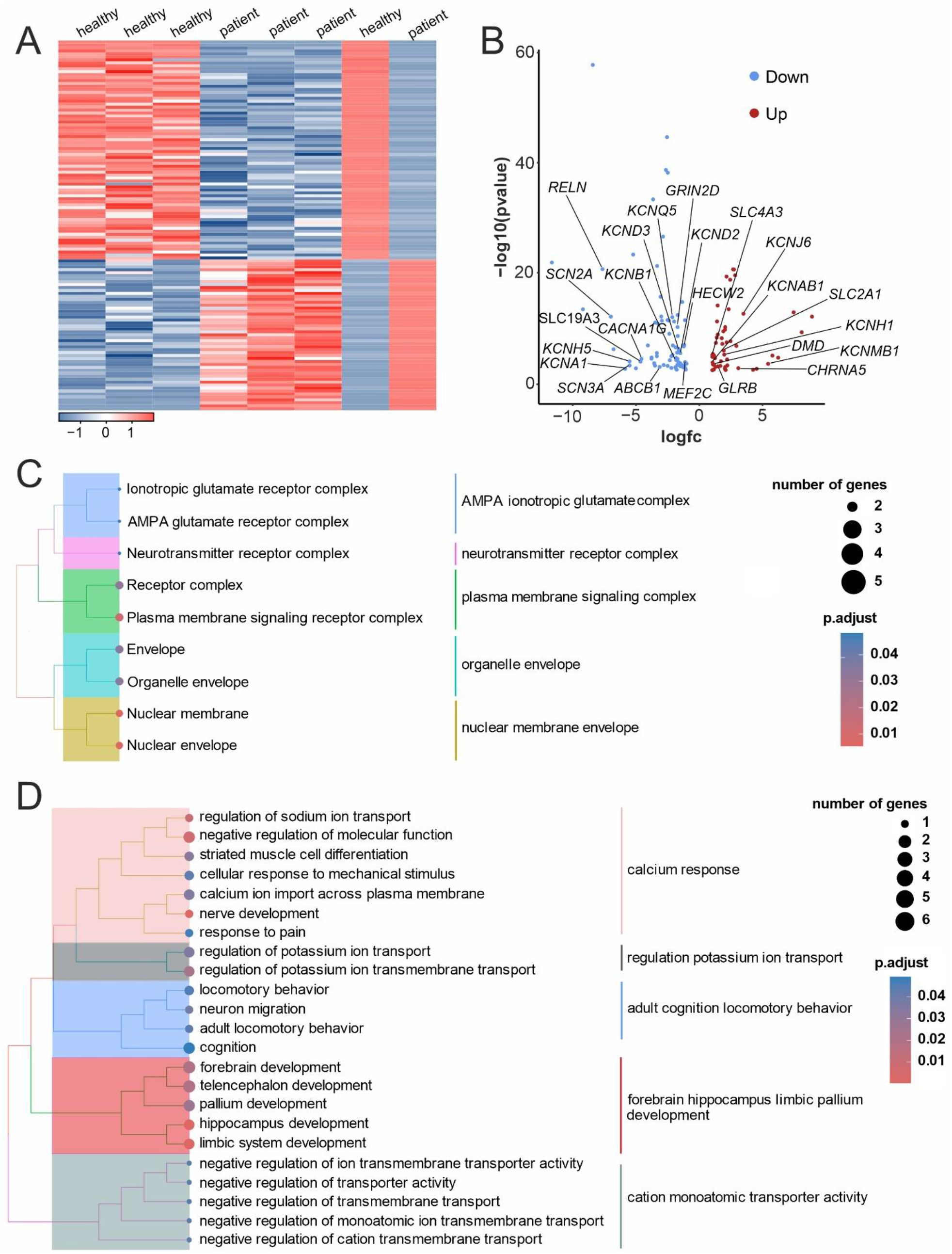
Transcriptome analysis of fibroblasts derived from patient and a healthy individuum. **(A)** DEGs of epilepsy-associated and transporter activity genes in patient derived fibroblasts are shown in heatmap analysis. Red color cells indicate upregulated genes and blue color cells indicate downregulated genes. **(B)** Volcano plot of DEGs shows the genes are common in epilepsy-associated genes and transporter activity related genes in patient-derived fibroblast. **(C)** Gene set enrichment (GSE) in total 125 DEGs shown in clusters. **(D)** GSE for “*cellular compartment*” analysis of differentially expressed genes. Only adjusted p-values less than 0.05 are shown. *n*=3 replicates for the analysis of healthy and USH2C fibroblasts.

We additionally performed GSEA on the DEGs which were detected in the patient fibroblasts. The *cellular compartment* GSEA showed that DEGs are related to “*AMPA ionotropic glutamate complex*”, “*neurotransmitter receptor complex*”, “*plasma membrane signaling complex*”, “*organelle envelope*” and “*nuclear membrane envelope*” (Figure 4C). GSEA of *biological function* showed that DEGs in patient fibroblast are associated with “*calcium response*”, “*regulation potassium ion transport*”, “*adult cognition locomotory behavior* “*forebrain, hippocampus, limbic and pallium development*” and “*cation monoatomic transporter activity*” (Figure 4D). GSEA of the “*biological function”* subclusters showed that DEGs in patient fibroblast are also related to “*nerve development and neuron migration”* as well as the “*development of the brain regions”*, such as the forebrain, hippocampus, and telencephalon (Figure 4D).

Altogether, Adgrv1 deficiency causes the dysregulation of genes related to glutamate homeostasis and epilepsy in mouse hippocampus and patient-derived cells.

### Glutamate uptake of primary brain astrocytes depends on Adgrv1 activation

Our omics data analysis suggested that ADGRV1 may play an important role in glutamate homeostasis in hippocampal astrocytes. To confirm the data at a functional level, we accessed glutamate uptake in primary astrocytes *in vitro* using a glutamate uptake assay introduced by (Mahmoud et al., 2019). Primary astrocytes were isolated from Adgrv1/del7TM and WT p0 mice and incubated with either 100 or 200 µM glutamate in a Ca^2+^ containing cell culture medium. Colorimetric analysis of the supernatants demonstrated the dose-dependent reduction in glutamate uptake by Adgrv1/del7TM astrocytes compared to WT controls (Figure 5A).

**Figure 5.**
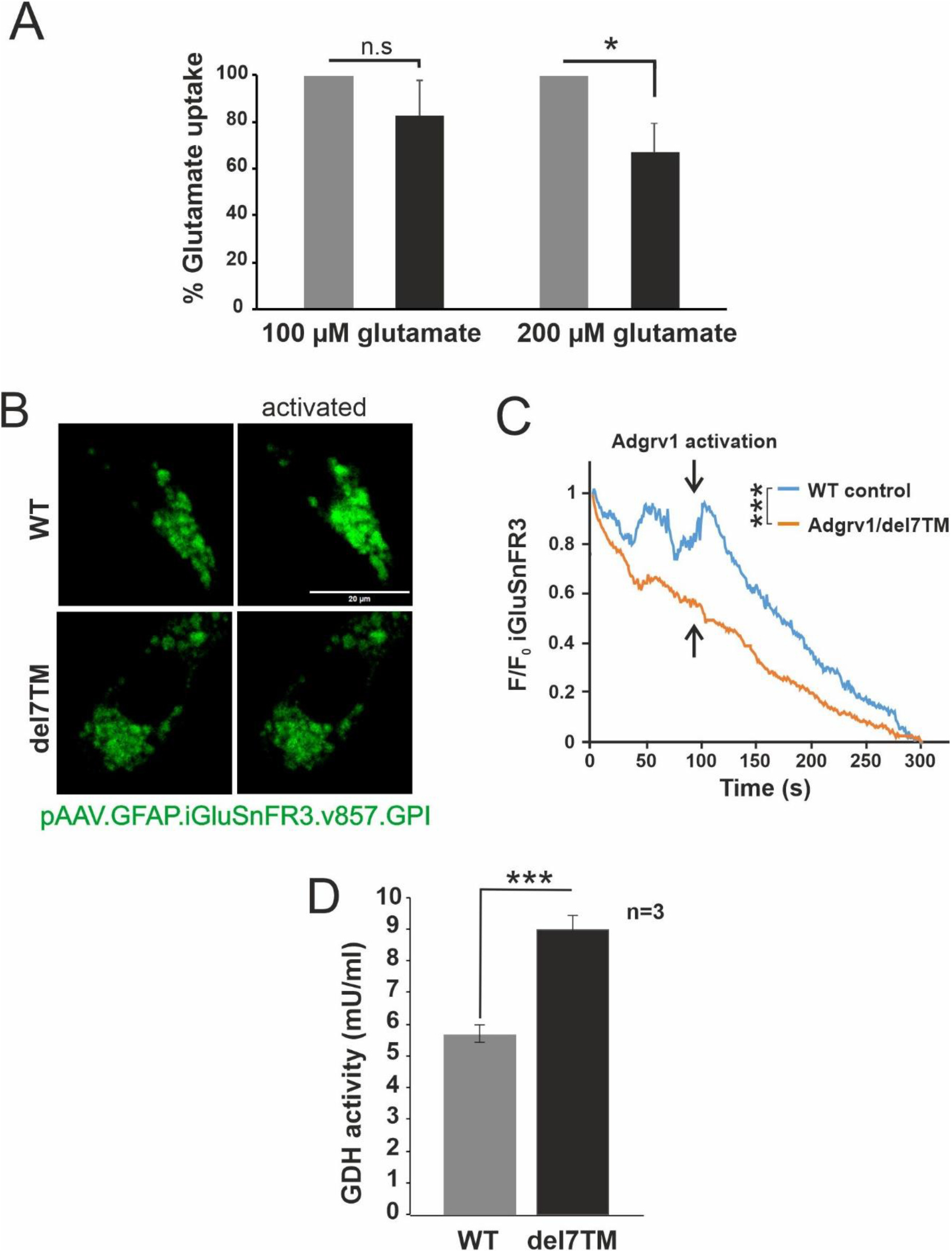
Glutamate uptake is mediated by Adgrv1 in primary astrocytes. (**A)** Glutamate uptake assay showing a dose-dependent reduction in Adgrv1/del7TM primary astrocytes. (**B)** Live-cell imaging of pAAV.GFAP.iGluSnFR3.v857.GPI transfected primary astrocytes derived from WT and Adgrv1/del7TM mice. Time-lapse image sequences of the fluorescent glutamate reporter were recorded with a sequence of 700 ms in a time course of 300 seconds with and without activation by the Statchel peptide. **(C)** Fluorescence intensity changes were calculated with F/F_0_ formula to obtain before and after application changes in fluorescence intensities. The yellow line indicates application of 1 mM “Stachel” peptide for receptor activation. F/F_0_ iGluSnFR3 intensity changes revealed an increase in WT astrocytes after Stachel peptide application, whereas there was no change observed in Adgrv1/del7TM astrocytes. (**D)** Glutamate dehydrogenase activity significantly increases in Adgrv1/del7TM astrocytes. For glutamate dehydrogenase activity assay 3 technical and 3 biological replicates were used for the quantification. For iGluSnFR3 intensity analysis *n*=7 (WT) and *n*=9 (Adgrv1/del7TM) in n=3 independent experiments were used. Data are represented as mean ± SD. Statistical evaluations were performed for bar plots using two-tailed Student’s t test and using Kruskal-Wallis test for iGluSnFr3 curves\**p* % 0.05, \*\**p* % 0.01, \*\*\**p* % 0.001. Scale bars: 20 µm.

To follow the uptake of glutamate from the media into the primary astrocytes, we used the genetically encoded glutamate reporter GFAP.iGluSnFR3.v857.GPI (Aggarwal et al., 2023). This glutamate reporter contains a glycosylphosphatidylinositol anchor that allows it to bind to glycosylphosphatidylinositols in biological membranes and thereby to monitor changes in glutamate concentration (Kinoshita and Fujita, 2016). In the first set of experiments, we confirmed that the glutamate reporter activity can be induced in both primary WT and Adgrv1/del7TM astrocytes by the addition of 10 mM glutamate to the culture medium (Figure S4). Next, we activated Adgrv1 using the “Stachel” of ADGRV1, an 11-amino acid peptide SVYAVYARTDN, which we had previously identified as the tethered agonist of the receptor (Knapp et al., 2022) to the culture medium of GFAP.iGluSnFR3.v857.GPI expressing primary astrocytes. Activation of Adgrv1 by the spike peptide led to an increase in reporter fluorescence in primary WT astrocytes, but not in astrocytes derived from Adgrv1-deficient Adgrv1/del7TM (Figure 5B, C). A control peptide, which consisted of a randomized amino acid sequence and should not activate Adgrv1, did not cause an increase in fluorescence in WT astrocytes (data not shown). Collectively, our data demonstrate that glutamate uptake into primary brain astrocytes depends on the activation of Adgrv1.

### The catabolism of internalized glutamate is increased in Adgrv1/del7TM hippocampal astrocytes

Alternative to its detoxification in the glutamate-glutamine cycle, glutamate can be also metabolized in astrocytes by TCA in mitochondria (Nissen et al., 2015). In the TCA, glutamate dehydrogenase (GDH) is the key enzyme, and its activity is a benchmark for glutamate metabolization. Measuring the GDH in cultured astrocytes with a colorimetric assay revealed an almost 1.5-fold increase in GDH activity in Adgrv1/del7TM astrocytes compared to WT astrocytes (Figure 5D).

### Expression of glutamine synthetase is reduced in hippocampus of Adgrv1/del7TM mice

Next, we analyzed the protein expression of key components of the glutamate-glutamine cycle in astrocytes of the hippocampal CA1 region: the glutamate transporter GLAST which imports glutamate from the extracellular space (Zhou et al., 2014) and the glutamine synthetase (GS) catalyzes the glutamine synthesis from glutamate in the cell (Papageorgiou et al., 2018) (Figure 6). We immunostained sections through the hippocampal CA1 region of Adgrv1/del7TM and WT mice for GS, GFAP, and DAPI (Figure 6A). Confocal microscopy demonstrated bright immunofluorescence of GS in GFAP-positive astrocytes of WT mice, and a greatly reduced fluorescence in astrocytes of Adgrv1/del7TM mice (Figure 6B). Quantification of anti-GS Western blots of hippocampus reveals a significant decrease in Adgrv1/del7TM mice (Figure 6C). In contrast, we observed only a minor and not significant increase in GLAST expression in the hippocampus of Adgrv1/del7TM neither by immunohistochemistry nor Western blotting (Figure 6D-F).

**Figure 6.**
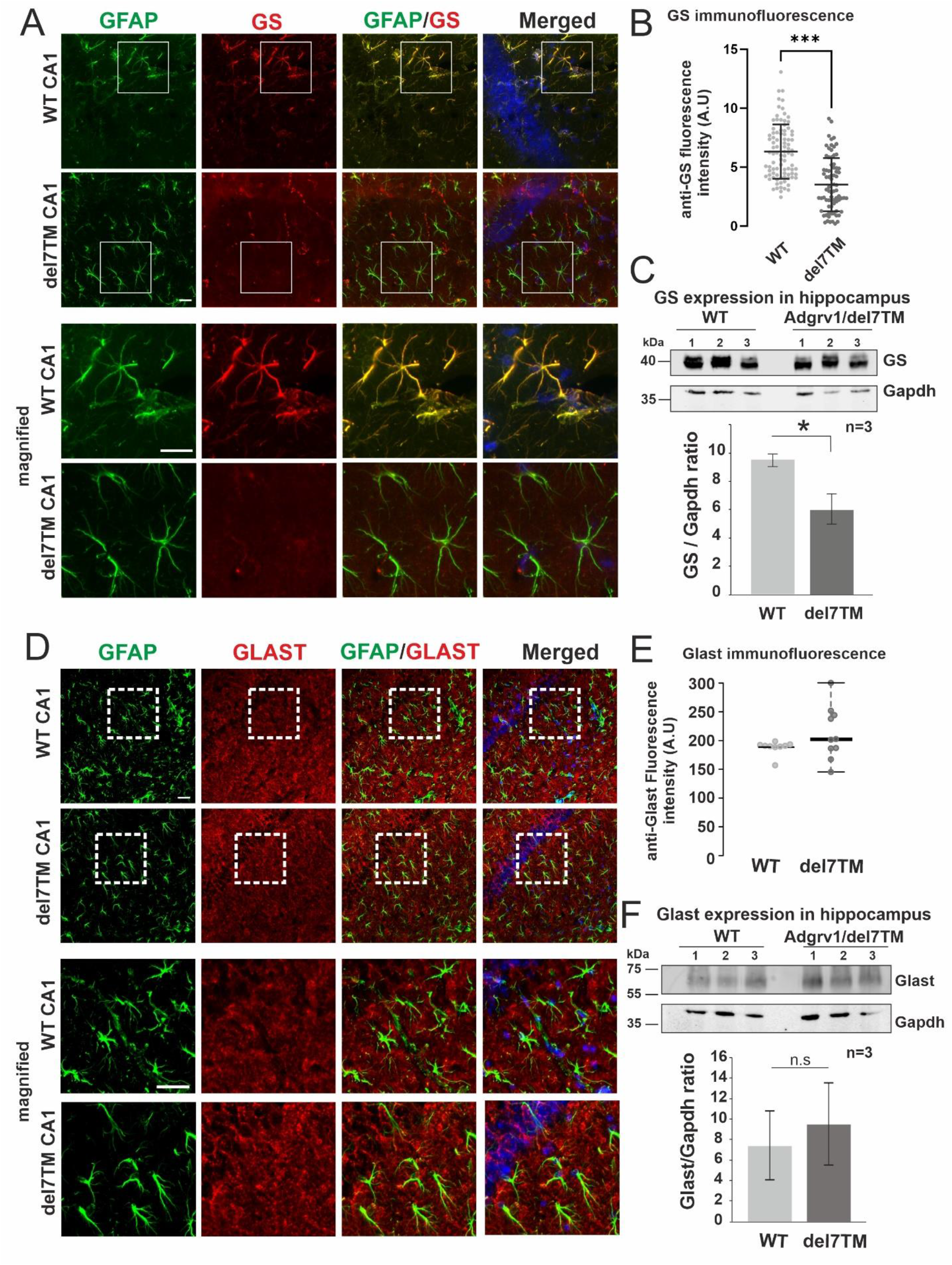
Expression analysis of glutamine synthetase (GS) and GLAST in astrocytes of the hippocampus of Adgrv1/del7TM and wild type control mice. **(A)** Double immunofluorescence staining for GS (red) and for the astrocyte marker GFAP (green) in sections through the CA1 region of the hippocampus counterstained for nuclear DNA by DAPI (blue). Merged images revealed localization and abundant expression of GS in GFAP-positive astrocytes in WT mice which is almost absent in Adgrv1/del7TM mice. **(B)** Quantification of anti-GS immunofluorescence intensity in in GFAP-positive astrocytes of hippocampal CA1 region reveals a high significant decrease in Adgrv1/del7TM mice. **(C)** Anti-GS Western blot analysis of hippocampal lysates demonstrates the significant decrease of GS protein expression in Adgrv1/del7TM mice. **(D)** Indirect immunofluorescence staining for GFAP (green) and glutamate transporter GLAST (red) counterstained by DAPI (blue) in the CA1 region of the hippocampus of Adgr1/del7TM (del7TM) and wild type (WT) mice. **(E)** Quantification of anti-GLAST immunofluorescence intensity in GFAP-positive astocytes of hippocampal CA1 region shows slight but not significant increase in Adgrv1/del7TM. *n*=3, number of sections quantified: 3 continuous sections per group. **(F)** Anti-GLAST Western blot analysis of hippocampal lysates of Adgrv1/del7TM and WT mice. Densitometry analysis related to Gapdh expression (A.U) showed tentative but an no upregulation of GLAST expression in Adgrv1/del7TM. *n*=3 animals and 3 continuous sections per group were used. Statistics: two-tailed Student’s t test, One way-ANOVA test; \**p*≤0.05, \*\**p*≤0.01, \*\*\**p*≤0.001. Scale bars: 20 µm. Magnified images Scale bars: 200 µm and 20 µm.

### Adgrv1 affects neurite morphogenesis in neurons

To investigate the effects of Adgrv1 in neurons, we analyzed single neuron cultures from brains of Adgrv1/del7TM and WT mice on day 1 *in vitro* (DIV1) and day 3 *in vitro* (DIV3) by staining for the neuronal marker MAP2 and with DAPI (Figures 7).

**Figure 7.**
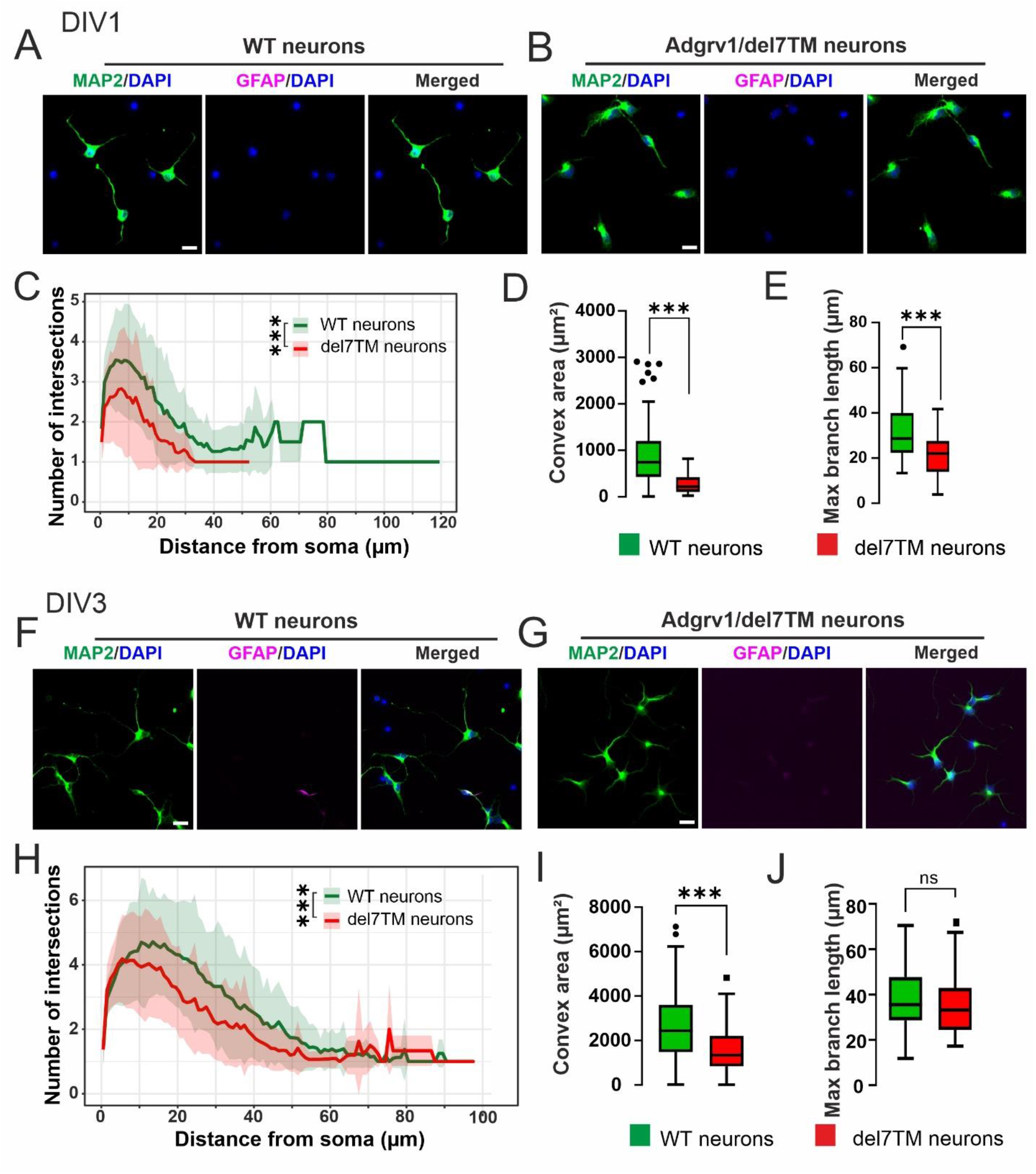
Morphometric characterization of primary neurons derived from WT and Adgrv1 mouse hippocampi. **(A, B)** Primary neurons isolated from the E18.5 WT and Adgrv1/del7TM mouse hippocampi were cultured for 1 day and stained with anti-MAP2 (green) and anti-GFAP (magenta) to visualize neurites of neurons and astrocytes, respectively. **(C)** Sholl analysis method applied for the quantification of intersection numbers of WT (green line) and Adgrv1/del7TM (red line) neuron cultures. A significant decrease in the number of neuronal intersections were observed in Adgrv1/del7TM neurons only cultures compared to the WT neuron cultures in day *in vitro* 1 (DIV1). **(D)** Analysis of Convex area showed a significant reduction in area of Adgrv1/del7TM neurons compared to WT neurons. **(E)** Maximum branch length of intersection in Adgrv1/del7TM neurons were also reduced. **(F)** WT and Adgrv1/del7TM neurons were cultured for 3 days in vitro (DIV 3). Neurons were identified with anti-MAP2 (green) and possible astrocyte contamination were detected with anti-GFAP (magenta) antibodies. **(H)** Quantification of **i**ntersection numbers by sholl analysis showed that Adgrv1/del7TM neurons have significantly less intersection numbers compared to WT neurons in DIV3. **(I)** Convex area quantification revealed that reduction in the area of Adgrv1/del7TM neurons persisted in DIV3 compared to WT neurons. **(G)** Maximum branch length of intersection analysis in DIV3 showed tentative decrease but no significant differences in between Adgrv1/del7TM and WT neurons. 60-70 cells in total for *n*=3 experiments. Statistics: One way-ANOVA test; \**p*≤0.05, \*\**p*≤0.01, \*\*\**p*≤0.001. Scale bar: 10 µm.

Morphological analysis of neurites in fluorescent microscopy images for neurite intersections (Sholl analysis), convex area, and maximal branch length revealed that all parameters were reduced in Adgrv1/del7TM neurons at both DIV1 and DIV3 (Figure 7C, H). All quantification analyses revealed significant differences (Figure 7D, I), except for the maximum branch length measurements in DIV3, which shows only a tendency in reduced length in Adgrv1/del7TM neurons (*p* value: 0.70) (Figure 7J). These findings suggest that Adgrv1 is not only important in astrocyte morphology (Figure 1C-F) but also has an additional significant influence on neuronal morphogenesis.

### Astrocyte Adgrv1 affects neurite morphogenesis

Astrocytes cooperate with neurons and support them in a variety of ways to maintain and nurture the neuronal microenvironment and help to control neuronal migration during development (Sidoryk-Wegrzynowicz et al., 2011). To investigate the effects of Adgrv1 in astrocytes on neurons, we analyzed cocultures of primary astrocyte and neurons from brains of Adgrv1/del7TM and WT mice at DIV1 and DIV3 and identified neurons and astrocytes with immunostaining of MAP2 and GFAP, respectively (Figures 8 and 9).

**Figure 8.**
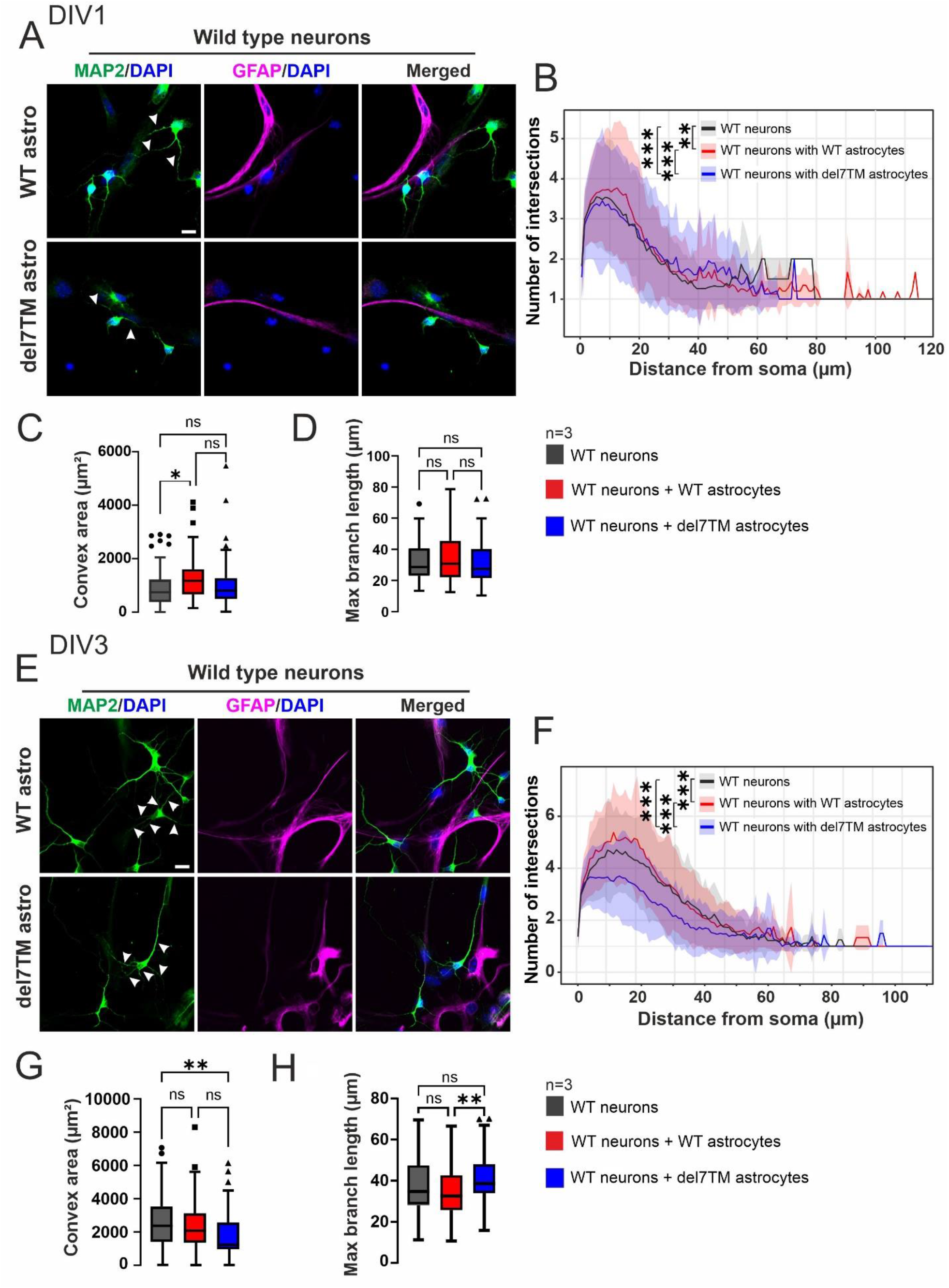
Morphometric characterization of primary neurons derived from WT mouse hippocampi co-cultured with WT and Adgrv1/del7TM astrocytes. **(A)** Primary neurons isolated from the WT E18.5 mouse hippocampus were cultured alone and together with WT or Adgrv1/del7TM astrocytes for DIV1. Anti-MAP2 (green) and anti-GFAP (magenta) were used for the visualization of neuronal neurites and astrocytes, respectively. Arrow heads indicate the neurites of primary neurons. (**B)** Sholl analysis revealed that co-culturing of WT neurons with WT astrocytes (red line) resulted in significant increase of intersection numbers of WT neurons. On contrary, co-culturing of WT neurons with Adgrv1/del7TM astrocytes (blue line) showed slight but significant decrease compared to WT neuron only cultures (black line). (**C)** Convex area analysis showed a notable increase in area of WT neurons co-cultured with WT astrocytes. However, no significant changes were observed in co-cultures with Adgrv1/del7TM astrocytes. (**D)** Maximum branch length of intersection in WT neurons did not change on both co-cultured together with WT and Adgrv1/del7TM astrocytes. (**E)** Immunofluorescence staining of neuronal neurites and astrocytes using Anti-MAP2 (green) and anti-GFAP (magenta), respectively in DIV3. (**F)** Total number of the intersections in WT neurons co-cultured with WT astrocytes (red line) showed the highest neurite numbers in DIV3 similar to DIV1 cultures. The analysis revealed that compared to WT neurons only cultures (black line) and WT neurons with WT astrocyte cultures (red line), co-culturing of WT neurons with Adgrv1/del7TM astrocytes (blue line) resulted in the lowest intersection numbers in DIV3 of culture. (**G)** Convex area quantification showed a significant decrease in WT neuron areas co-cultured with Adgrv1/del7TM astrocytes in DIV3. However no significant changes were observed in the WT neuron area when co-cultered with WT astrocytes. **(G)** Maximum branch length of intersections in WT neurons did not change in both co-cultures together with WT and Adgrv1/del7TM astrocytes in DIV3. However, analysis revealed that WT neurons with WT astrocytes co-cultures have significantly smaller areas compared to WT neurons with Adgrv1/del7TM astrocytes. 60-70 cells in total for *n*=3 experiments. Statistics: One way-ANOVA test; \**p*≤0.05, \*\**p*≤0.01, \*\*\**p*≤0.001. Scale bar: 10 µm.

**Figure 9.**
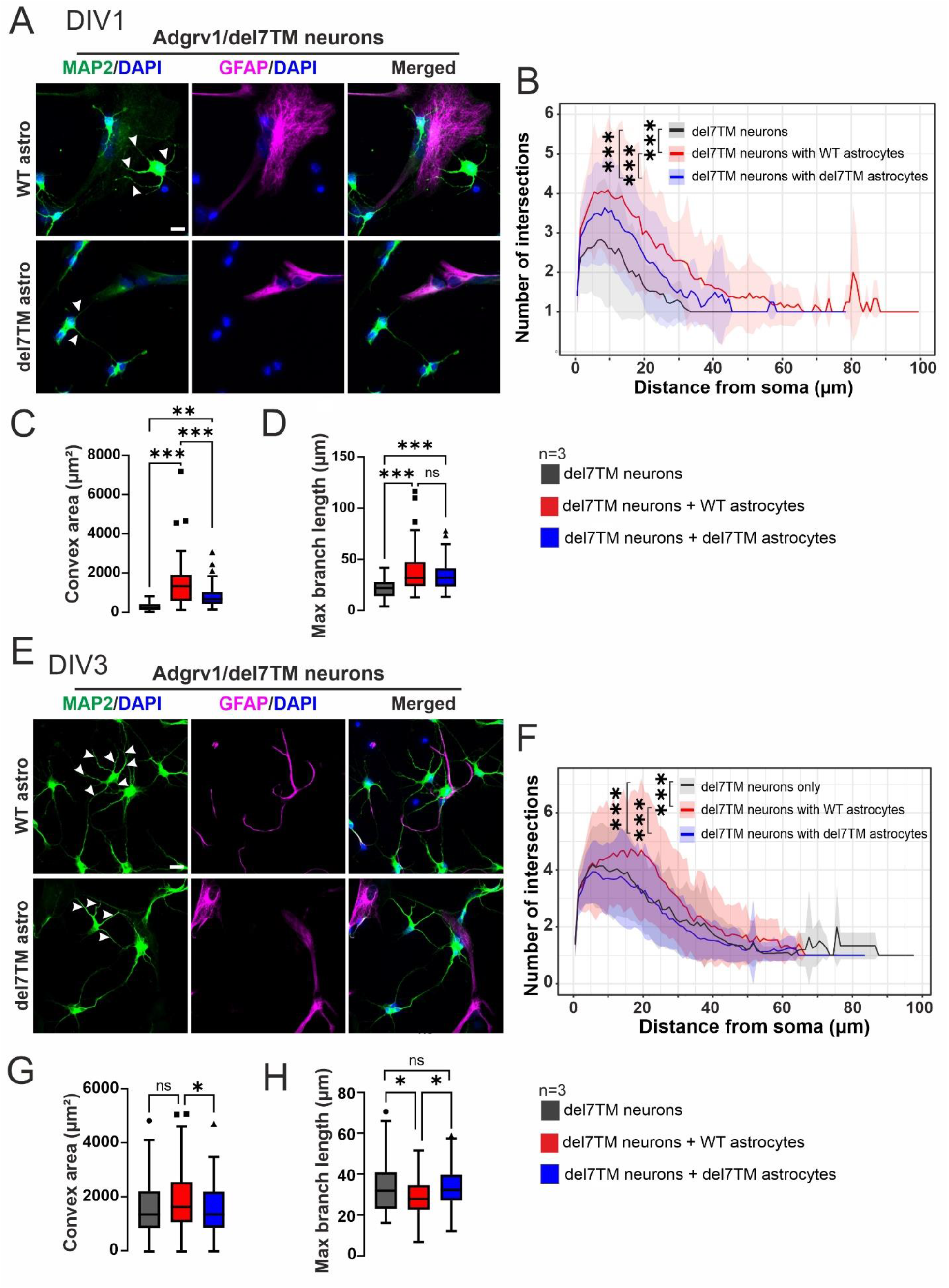
Morphometric characterization of primary neurons derived from Adgrv1/del7TM mouse hippocampi co-cultured with WT and Adgrv1/del7TM astrocytes. **(A)** Primary neurons isolated from the Adgrv1/del7TM E18.5 mouse hippocampus were cultured alone or together with WT or Adgrv1/del7TM astrocytes for DIV1. Anti-MAP2 (green) and anti-GFAP (magenta) were used for the visualization of neuronal neurites and astrocytes, respectively. Arrow heads indicate the neurites of neurons. (B) Sholl analysis of Adgrv1/del7TM neurons only cultures (black line) revealed the least intersection numbers. Co-culturing of the Adgrv1/del7TM neurons with WT (red line) or Adgrv1/del7TM (blue line) resulted in significant increase in intersection numbers. The most prominent increase in intersection numbers was observed in Adgrv1/del7TM neurons co-cultured with WT astrocytes. (**C)** Co-cultures of Adgrv1/del7TM neurons with WT or Adgrv1/del7TM astrocytes significantly increased the area of Adgrv1/del7TM neurons. The most prominent increases in the Adgrv1/del7TM neuron areas were observed in co-cultures with WT astrocytes. (**D)** Maximum branch length of intersection in Adgrv1/del7TM neurons significantly increased in both co-cultured together with WT and Adgrv1/del7TM astrocytes in DIV1. (**E)** Double immunofluorescence staining of neuronal neurites and astrocytes using Anti-MAP2 (green) and anti-GFAP (magenta), respectively in DIV3. (**F)** The highest intersection numbers were observed in Adgrv1/del7TM neurons when co-cultured with WT astrocytes (red line). Adgrv1/del7TM neurons co-cultured with Adgrv1/del7TM astrocytes (blue line) showed a significantly lowest intersection numbers compared to both Adgrv1/del7TM neuron only cultures (black line) and Adgrv1/del7TM neurons/WT astrocyte co-cultures in DIV3. (**G)** Co-culturing the Adgrv1/delTM neurons with both WT and Adgrv1/del7TM astrocytes did not change the area of Adgrv1/del7TM neurons. However, a significant reduction observed in Adgrv1/del7TM neuron and Adgrv1/del7TM astrocyte co-cultures compared to co-cultures with WT astrocytes in DIV3. **(G)** Maximum branch length of intersections in Adgrv1/del7TM neurons co-cultured with WT astrocytes showed significant reduction compared to Adgrv1/del7TM neuron only and Adgrv1/del7TM neuron co-cultured with Adgrv1/del7TM astrocytes in DIV3. 60-70 cells in total for *n*=3 experiments. Statistics: One way-ANOVA test; \**p*≤0.05, \*\**p*≤0.01, \*\*\**p*≤0.001. Scale bar: 10 µm.

Sholl analysis of the fluorescence images indicated a significant increase in the number of neurite intersections in WT neurons when co-cultured with WT astrocytes at both DIV1 and DIV3, indicating a synergistic effect on neuronal development (Figure 8B, F). In contrast, co-culturing WT neurons with Adgrv1/del7TM astrocytes led to fewer neurite intersections at both stages, with the most pronounced effect at DIV3 (Figure 8B, F). Furthermore, the convex area of neurites in WT neurons was significantly increased by WT astrocytes at DIV1 but not at DIV3, suggesting a temporal influence on neurite morphology (Figure 8C, G). Interestingly, DIV3 cultures with Adgrv1/del7TM astrocytes exhibited a decrease in convex area compared to WT neuron cultures alone, highlighting the impact of Adgrv1 deficiency on neurite development (Figure 8C, G). However, the maximum branch length analysis revealed no significant differences in WT neurons when co-cultured with either WT or Adgrv1/del7TM astrocytes at DIV1 and DIV3 (Figure 8D, H).

Next, we explored the role of astrocyte Adgrv1 expression on Adgrv1-deficient neurons. Sholl analysis revealed a substantial increase in neurite numbers in the presence of WT astrocytes at both DIV1 and DIV3 (Figure 9B, F). Additionally, WT astrocytes significantly increased the convex area of neurites in WT neurons at DIV1, though not at DIV3, indicating a dynamic interplay between astrocytic Adgrv1 expression and neurite morphology (Figures 9C, G). Moreover, maximum branch length analysis of Adgrv1/del7TM neurons showed an initial increase at DIV1 followed by a significant decrease at DIV3 in the presence of WT astrocytes (Figure 9D, H). However, Adgrv1/del7TM astrocytes showed an opposite effect on Adgrv1/del7TM neurons, namely an increase of the number of intersection numbers and of convex area of neurites at DIV1 (Figure 9B, C) but inducing a significant decrease at DIV3 cultures (Figure 9F, G).

In summary, Adgrv1 in astrocytes supports the beneficial role of astrocytes in neurite morphogenesis during neuronal development and the dysfunctions of Adgrv1-deficient astrocytes exacerbate defective neurite morphogenesis in Adgrv1-deficient neurons.

### Adgrv1 functions in the astrocyte-neuron crosstalk affecting synaptogenesis

Astrocytes promote synapse formation in a variety of ways (Baldwin and Eroglu, 2017). To determine whether Adgrv1 in astrocytes plays a role in synaptogenesis in neurons, we isolated primary neurons and astrocytes derived from the hippocampus of WT or Adgrv1/del7TM mice and cultured primary neurons alone or together with primary astrocytes, respectively. In DIV14 cultures, we immunostained neurons for homer, a post synaptic density (PSD) marker for excitatory synapses, gephyrin as a PSD marker for inhibitory synapses, and the neurite marker MAP2. This triple labelling allowed us to determine the number of PSD per 100 µm neurite length ∼ (synaptic density) and sizes of the PSD puncta (∼ synaptic strength) (Holler et al., 2021), and to distinguish between the excitatory and inhibitory synapses in neurites (Figure 10A).

**Figure 10.**
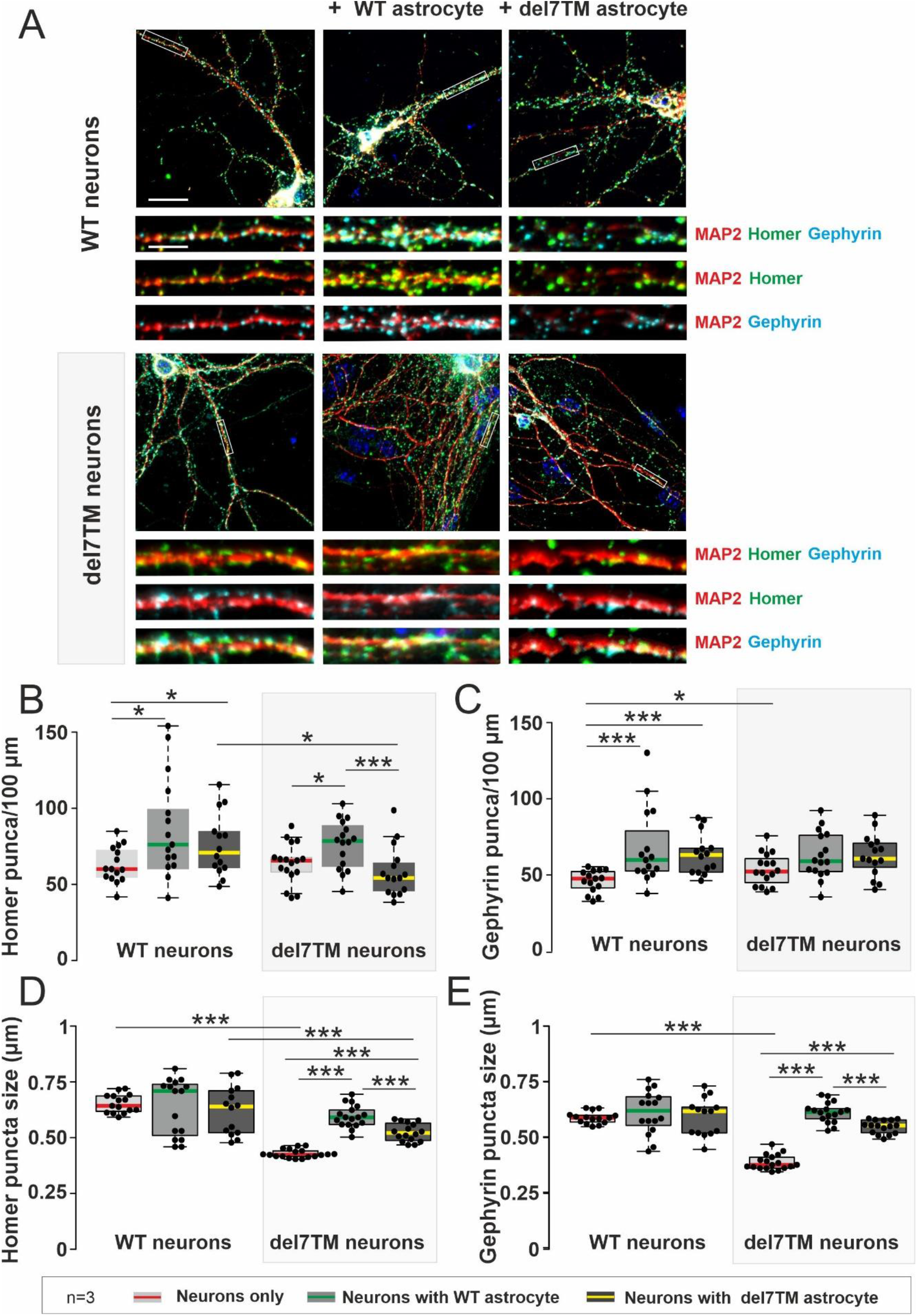
Co-culturing of WT and Adgrv1/del7TM neurons with WT astrocytes increases synaptic numbers and sizes. **(A)** Hippocampal neurons from WT and Adgrv1/del7TM mice were immunolabelled with anti-Homer (Green), anti-Gephyrin (Cyan) and anti-MAP2 (red). Nucleus was counter-stained with DAPI. **(B)** Quantification of excitatory post-synaptic puncta revealed that similar Homer puncta in WT and Adgrv1/del7TM neuron cultures. WT astrocyte and neuron co-cultures increased the Homer puncta number in both neuron cultures. The lowest homer puncta number was observed in Adgrv1/del7TM astrocyte-neuron co-cultures. **(C)** Quantification of inhibitory post-synaptic puncta using Gephyrin staining showed significantly more puncta in Adgrv1/del7TM neuron only cultures compared to WT neuron cultures. While astrocyte co-cultures increased the numbers of inhibitory synaptic puncta in WT neuron co-cultures, no differences observed in Adgrv1/del7TM neuron co-cultures. **(D)** Excitatory post-synaptic puncta size did not change in WT neuron co-cultures with WT or Adgrv1/del7TM astrocytes. Adgrv1/del7TM neurons showed significantly smaller size of Homer puncta. Smaller size observed in Adgrv1/del7TM neurons were increased by co-culturing with WT or Adgrv1/del7TM astrocytes. **(E)** Similar to excitatory synapses quantification, there was a significant decrease in inhibitory synaptic puncta in Adgrv1/del7TM neurons. Highest puncta sizes were observed in neurons with WT astrocyte co-cultures. Scale bar: 10 and 20 µm. 15-16 cells in total for *n*=3 experiments were used, and statistical evaluation was performed using Student’s t-test analysis: \**p*≤0.05, \*\**p*≤0.01, \*\*\**p*≤0.001.

In neuron-only cultures of WT or Adgrv1/del7TM mice, there were no differences in the density of excitatory PSDs (homer puncta) on neurites while the density of inhibitory PSDs (gephyrin puncta) was slightly increased (Figure 10B, C). In co-cultures of WT neurons with astrocytes from either WT and Adgrv1/del7TM mice, the density of synaptic puncta was significantly increased in both excitatory and inhibitory synapses (Figure 10B, C) compared to neuron-only cultures. Co-culturing of Adgrv1/del7TM neurons with both WT and Adgrv1/del7TM astrocytes, respectively, did not affect the inhibitory PSD puncta density on neurites (Figure 10C). However, excitatory PSD puncta density on Adgrv1/del7TM neurites increased to the level of WT neurites in co-cultures with WT astrocytes but decreased in co-cultures with Adgrv1/del7TM astrocytes (Figure 10B). Our data suggest that Adgrv1 in astrocytes controls the frequency of excitatory synapses on neurites but has only a modest effect on the frequency of inhibitory synapses. This may lead to an imbalance of excitatory-inhibitory (E/I) synapses which has been previously correlated with neurological diseases such as epilepsy (Bonansco and Fuenzalida, 2016).

Next, we measured and quantified the size of the synaptic puncta stained for homer and gephyrin, a feature previously shown to be related with synaptic strength (Holler et al., 2021) (Figure 10D, E). In WT neurons the puncta size for both excitatory and inhibitory PSDs was not affected by co-cultured WT and Adgrv1/del7TM astrocytes. However, in neuron-only cultures, the sizes of the excitatory and inhibitory PSDs were significantly decreased (*p*-value 1.29987e^-19^ and 1.79652e^-18^, respectively) in Adgrv1/del7TM compared to WT control mice. This was compensated in co-cultures with Adgrv1/del7TM or WT astrocytes but were more prominent by WT astrocytes (Figure 10D, E).

In summary, our data demonstrate that Adgrv1 expression in neurons significantly affects the size of PSDs which correlates with a reduced synaptic strength in Adgrv1-deficient neurons. Furthermore, our data show that the synaptic strength in Adgrv1-deficient neurites increases by the interaction with astrocytes and that Adgrv1 in astrocytes can potentiate this effect.

## DISCUSSION

The high expression of ADGRV1 in astrocytes is indicative of its major importance for astrocyte functions which was confirmed by our omics data: Using affinity proteomics, we identified nearly 300 proteins enriched in the proteome of human and mouse astrocytes (Batiuk et al., 2020) as potential interacting proteins of ADGRV1. In addition, transcriptome analyses of mouse hippocampus and human cells deficient for ADGRV1 revealed that numerous genes encoding for proteins with important functions in astrocytes were differentially expressed.

Furthermore, our omics data analysis suggests that ADGRV1 is associated with signaling pathways that are highly specific for astrocytes and essential for their proper function. One of the major functions of astrocytes in the brain is the rapid clearance of excitatory neurotransmitter glutamate from the synaptic cleft after its release from the pre-synapse to prevent glutamate neurotoxicity (Armbruster et al., 2016; Langer et al., 2017). Strikingly, we identified the glutamate transporters EAAT3 and EAAT1 (GLAST1) as potential interacting partners of ADGRV1 (Zhou et al., 2014; Piepgras et al., 2022). Although we did not find altered protein expression of glutamate transporters in Adgrv1-deficient cells, the interaction of ADGRV1 with these core proteins of the glutamate uptake machinery suggests a role in the regulation of glutamate uptake processes in astrocytes. Indeed, in *in vitro* glutamate uptake assays and by live cell imaging using a glutamate reporter, we found consistently reduced glutamate uptake in primary astrocytes deficient for Adgrv1 confirming that ADGRV1 participates in glutamate uptake in astrocytes. Interestingly, glutamate uptake by astrocytes could be triggered by the synthetic activator mimicking the tethered agonist peptide, (Knapp et al., 2022). This stimulation suggests that glutamate uptake is controlled by the activation of the CTF receptor part of ADGRV1 and is not due to its cell adhesion function. Failure in glutamate uptake by astrocytes and thereby of the clearance of glutamate form the extracellular environment can lead to glutamate toxification in the brain and lead to glia and neuron cell death (Kritis et al., 2015). The toxicity of excess glutamate in the extracellular *milieu* is probably the cause for the astrocyte depletion found in in the hippocampus Adgrv1/del7TM mice.

After uptake by astrocytes, glutamate can be converted into the non-toxic essential amino acid glutamine catalyzed by GS as part of the glutamate-glutamine cycle between astrocytes and neurons (Papageorgiou et al., 2018). Alternatively, glutamate can be also metabolized in astrocytes by the TCA cycle which operates inside the mitochondria (Rose et al., 2020). A balanced interaction of the two pathways is vital for glutamate homeostasis in astrocytes and globally in the brain (Robinson et al., 2020). The identification of potential interacting proteins of ADGRV1 and DEGs in Adgrv1/del7TM mice related to both pathways collectively support a vital role of ADGRV1 in glutamate homeostasis. This role is further supported by the dysregulation of two key enzymes of the two pathways in Adgrv1-deficient astrocytes, namely an increase of the activity of the glutamate dehydrogenase (GDH) in the TCA and the glutamate synthetase (GS) a pivotal enzyme in the glutamate-glutamine cycle.

Interestingly, the increase of the activity of the TCA enzyme GDH is accompanied in Adgrv1-deficient astrocytes by a drastic reduction in the expression of the GS protein, a key enzyme in the glutamate-glutamine cycle. The downregulation of GS in ADGRV1-deficient hippocampal astrocytes is likely due to deficient glutamate uptake, as described in previous studies, leading to accumulation of toxic intracellular glutamate in astrocytes (Trabelsi et al., 2017). Intriguingly, a decrease of GS expression accompanied with an increase of GDH activity was previously described in patients with myoclonic absence epilepsy with sclerotic hippocampus (D.M. et al., 2005; Bahi-Buisson et al., 2008; Chan et al., 2019) a clinical condition that also occurs in ADGRV1-associated epilepsy (Leng et al., 2022). Overall, our results suggest that ADGRV1 is significantly involved in the control of glutamate homeostasis, which is imbalanced by defects in ADGRV1, which is proven to be disease relevant (Onaolapo and Onaolapo, 2020).

In the brain, ADGRV1 is strongly expressed in neurons, but highest in astrocytes (McMillan and White, 2010) Here, we observed that deficiency of ADGRV1 leads to decreased abundance and altered morphology of astrocytes in the CA1 region of the hippocampus, most likely due to an imbalanced glutamate homeostasis. Corresponding changes in astrocyte morphology and number have been observed in diseases in which astrocytes are known to be involved in the pathogenesis such as acute brain trauma and chronic neuropathies, *e.g*., Alzheimer’s disease, but also in the development of epilepsy (Wu et al., 2021; Hayashi et al., 2022; de Sousa et al., 2023). The increase in branching or convex hull area of astrocytes is thought to be a consequence of the reduced number of cells in the diseased brain due to their exploratory attempts to contact the remaining neurons.

In the present study, we demonstrate that ADGRV1 is not only important for the functions of astrocytes but also of neurons. In developing neurons, the lack of ADGRV1 compromises neurite synaptogenesis. This is consistent with previously found interactions of ADGRV1 with numerous components of the pre- and post-synapse and with molecules important for synaptic signaling and neurotransmitter vesicle cycle (Knapp et al., 2019). The previously identified interactions of ADGRV1 with other synaptic molecules such as latrophillin 2 (ADGRL2) another aGPCR (Anderson et al., 2017), might be vital for synaptic function and synaptogenesis. This hypothesis supports our finding that expression of Adgrv1 in astrocytes is crucial for the size of excitatory and inhibitory synapses and thereby for synaptic strength (Chung et al., 2015; Gürth et al., 2020). Interestingly, the presence of Adgrv1 in astrocytes influences only the abundance of excitatory synapses on neurites, but not of inhibitory synapses. This possibly leads to an imbalance of excitatory-inhibitory (E/I) synapses previously correlated with neurological diseases such as epilepsy (Bonansco and Fuenzalida, 2016).

In addition, we found that neurite branching was significantly reduced in ADGRV1-deficient neurons. This reduction is consistent with the identification of DEGs in patient cells with mutations in *ADGRV1* that encode proteins related to neuronal development and neuronal migration as well as the development of brain regions. These results demonstrate a possible role of ADGRV1 in neurite outgrowth during neurogenesis previously proposed (Knapp et al., 2022).

The molecular crosstalk between astrocytes and neurons is crucial for the correct development, function, and maintenance of the proper health of the CNS (Allen and Eroglu, 2017; Perez-Catalan et al., 2021). Here, we studied the contribution of ADGRV1 in astrocyte-neuron communication in co-cultures of primary cells from the mouse hippocampus. We found that developing primary WT neurons were supported in their development by the presence WT astrocytes but not by Adgrv1-deficent astrocytes. In addition, the defective development of Adgrv1/del7TM primary neurons was rescued by the presence of WT primary astrocytes but was potentiated by Adgrv1-deficent astrocytes. Our findings suggest the bidirectional interaction of ADGRV1 in astrocyte-neuron communication supportive for the correct development of neurons.

Pathogenic variants of *ADGRV1* cause quite distinct diseases such as human Usher syndrome type 2 leading to hereditary deaf blindness (Weston et al., 2004) and various forms of epilepsy in human and rodents (McMillan and White, 2010; Wang et al., 2015; Myers et al., 2018; Liu et al., 2020; Leng et al., 2022; Zhou et al., 2022). It can be assumed that the loss of the fibrous membrane links formed by the extracellular domain of ADGRV1 cause crucial defects in the sensory cells of the inner ear and eye leading to USH2C (Lefèvre et al., 2008; Maerker et al., 2008). It is possible, but no current evidence suggests, that dysfunctions of ADGRV1 in astrocytes may also contribute to the USH disease in eye and the inner ear links.

The data presented in this paper provides first insights into the molecular and cellular basis underlying epilepsy associated with mutations in *ADGRV1*. Present omics data identified molecules as potential interaction partners of ADGRV1 related to glutamate homeostasis (see above) have been previously also associated with epilepsy (Lechan and Fekete, 2006; Poduri et al., 2013; Falk et al., 2014; Saleh et al., 2020; Sachs et al., 2023). The absence of ADGRV1 from these protein complexes may also result in pathways altered in epilepsy. This is consistent with our finding that DEGs found in the hippocampus of Adgrv1/delTM7 mice and in patient-derived cells with ADGRV1 deficiency have previously been associated with epilepsy. Interestingly, the morphological and physiological alterations in astrocytes were pronounced especially in the CA1 of the hippocampus, a region which is vulnerable to glutamate toxicity (Ouyang et al., 2007), where recently astrocyte loss was related to early epileptogenesis (Wu et al., 2021). Cumulatively, our data support the hypothesis that the molecular origin of ADGRV1-associated epilepsy lies in the dysfunction of astrocytes in the hippocampus due to impaired glutamate homeostasis and its consequences caused by ADGRV1 deficits.

## Conclusion

We show here that ADGRV1 is crucial for the morphology and physiology of hippocampal astrocytes. ADGRV1 deficiency imbalances the glutamate homeostasis possibly leading to glutamate toxification in the cell and neuronal tissue. The resulting consequences presumably lead to impaired neuronal development and to impaired communication between astrocytes and neurons in the brain. These molecular dysfunctions of ADGRV1-deficent astrocytes provide first clues to understanding the pathomechanisms in epilepsy associated with mutations in *ADGRV1* and will be essential in the development of future therapies. Future studies aimed at the physiology of receptor complexes in astrocytes and neurons related to ADGRV1 and the pathways downstream to activated ADGRV1 could assist to defining the precise mechanisms of ADGRV1 functions in the CNS in health and disease. This should also provide further targets for therapies and offer prospects for the future treatment and cure of ADGRV1-related epilepsy.

## Acknowledgements

We thank Ulrike Maas, and Yvonne Kerner for excellent technical assistance, and Drs. Filip Maciag for critical discussion of the data. We also thank Kevin Peter Langenberg and Selina Kröll for their support in the analysis of neuron morphology. We thank Dr. Erwin van Wijk for kindly providing patient fibroblasts for the present study.

## Supplementary Materials

**Table S1.** Biological function analysis of astrocyte related proteins in ADGRV1 TAPs and protein annotations. Subtitle 1 shows the astrocytes enriched proteins and their annotations. Subtitle 2 shows biological functions of astrocytes enriched proteins. Subtitle 3 shows transporter activity and L-glutamate import related proteins.

**Table S2.** DEGs in WT and Adgrv1/del7TM mouse hippocampi.

**Table S3.** The result of RNAseq analysis in fibroblast derived from patient. Subtitle 1 shows DEGs in patient-derived fibroblast. Subtitle 2 shows transporter activity genes found in patient derived fibroblasts. Subtitle 3 shows epilepsy associated genes DEGs found in patient fibroblasts. Subtitle 4 indicates common DEGs in transporter activity and epilepsy associated genes. Subtitle 5 shows total 126 DEGs found in transporter activity and epilepsy associated genes.

## Institutional Review Board Statement

The use of mice in research was approved by the German regulation authority for the use of animals in research, the district administration Mainz-Bingen, 8341a/177-5865-§11 ZVTE, 30.04.2014.

## Data availability statement

Full Western blots presented in the study are included in the article/Supplementary Material. TAP data have been deposited to the ProteomeXchange Consortium via the PRIDE partner repository with the dataset identifier PXD042629. Codes for the RNA-sequencing analysis can be found at https://github.com/LabWolfrum.

## Author contributions

All authors contributed to the article and approved the submitted version. B.E.G. conducted most of the experiments, analyzed tandem affinity purification data sets, and prepared all the figures for publication. J.L. helped with the isolation of hippocampus RNA samples and contributed to TAP data analysis. M.Z. performed the analysis of RNA sequencing. B.E.G. and U.W. conceptualized the study and wrote the manuscript. K.N.-W. supervised M.Z., contributed with patient-derived fibroblast cultures, discussing data and writing.

## Supplemental Figures

**Figure S1.**
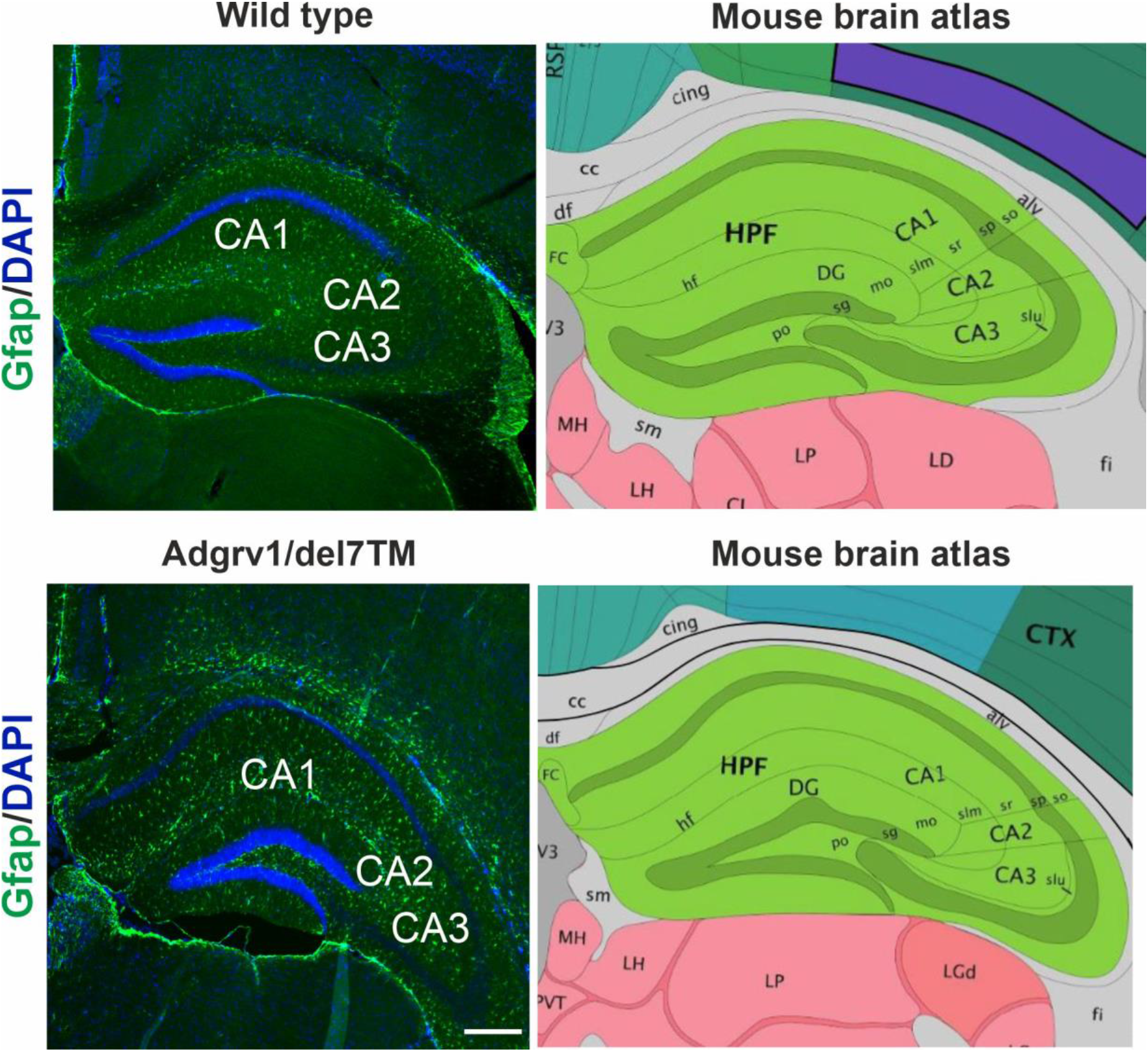
Reference coronal brain sections from mouse brain atlas for the identification of hippocampus subregions. The subregions of the mouse hippocampus sections were identified using the Allen mouse brain atlas (https://mouse.brain-map.org/) during the image analysis. Scale bar: 25 µm.

**Figure S2.**
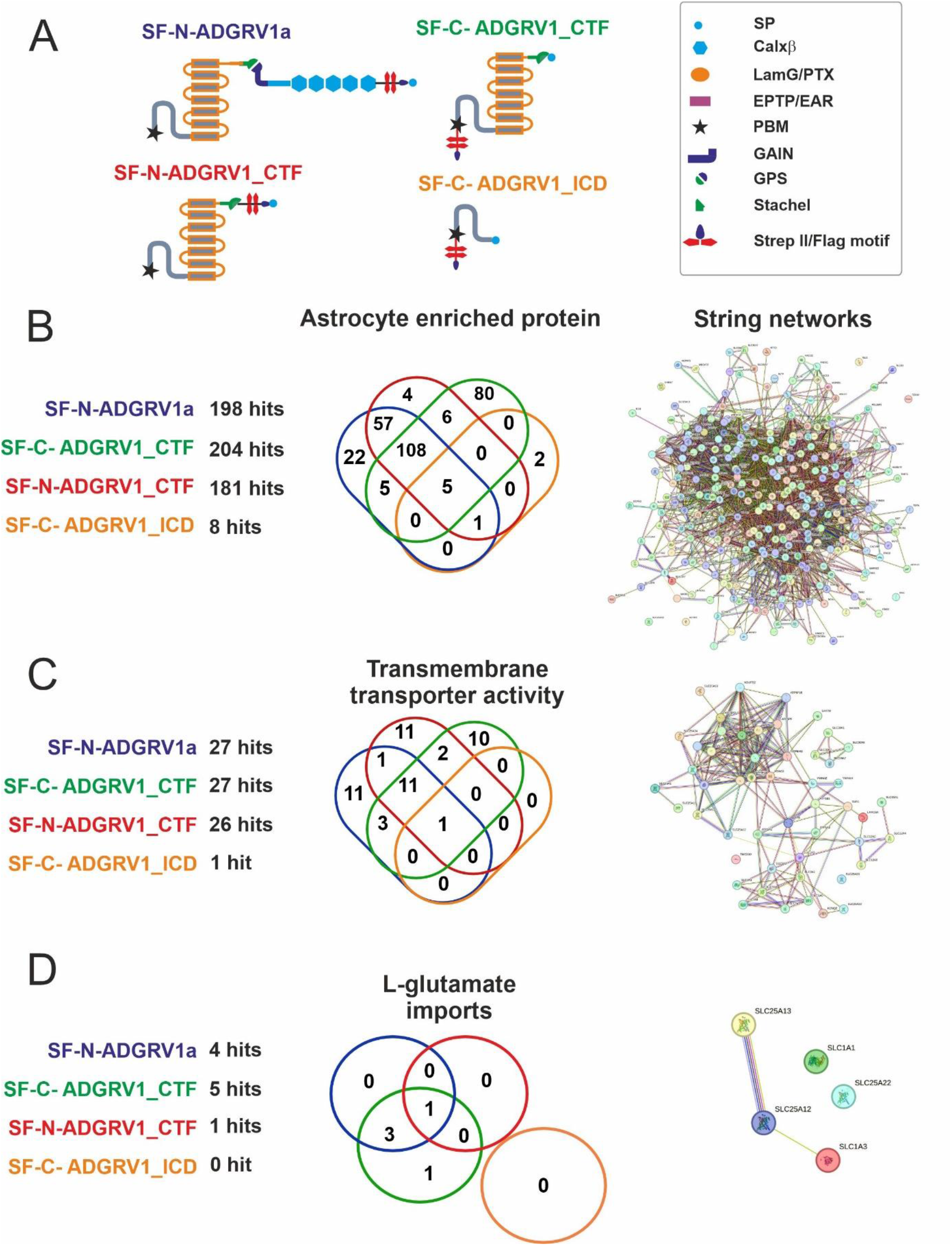
TAP analysis reveals a complex protein network related to ADGRV1 protein. **(A)** Different Strep II/FLAG (SF)-tagged ADGRV1 constructs used as a prey in Tanden affinity purification (TAP) to revealing potential interaction partners of ADGRV1 using HEK293T cells. ADGRV1 constructs were tagged from C or N terminals to eliminate false binding partners and changes in receptor structure. (**B**) GO term analysis revealed astrocyte enriched protein in ADGRV1 TAP analysis. The Venn-diagram shows a high overlap in astrocyte enriched proteins in 4 different ADGRV1 constructs, and string network analysis shows interactions. (**C**) Transmembrane transporter activity related proteins highly overlapped in SF-N-ADGRV1a, SF-N-ADGRV1_CTF and SF-C-ADGRV1 preys. (**D**) L-glutamate import related proteins have enriched in SF-N-ADGRV1a and SF-C-ADGRV1 preys.

**Figure S3.**
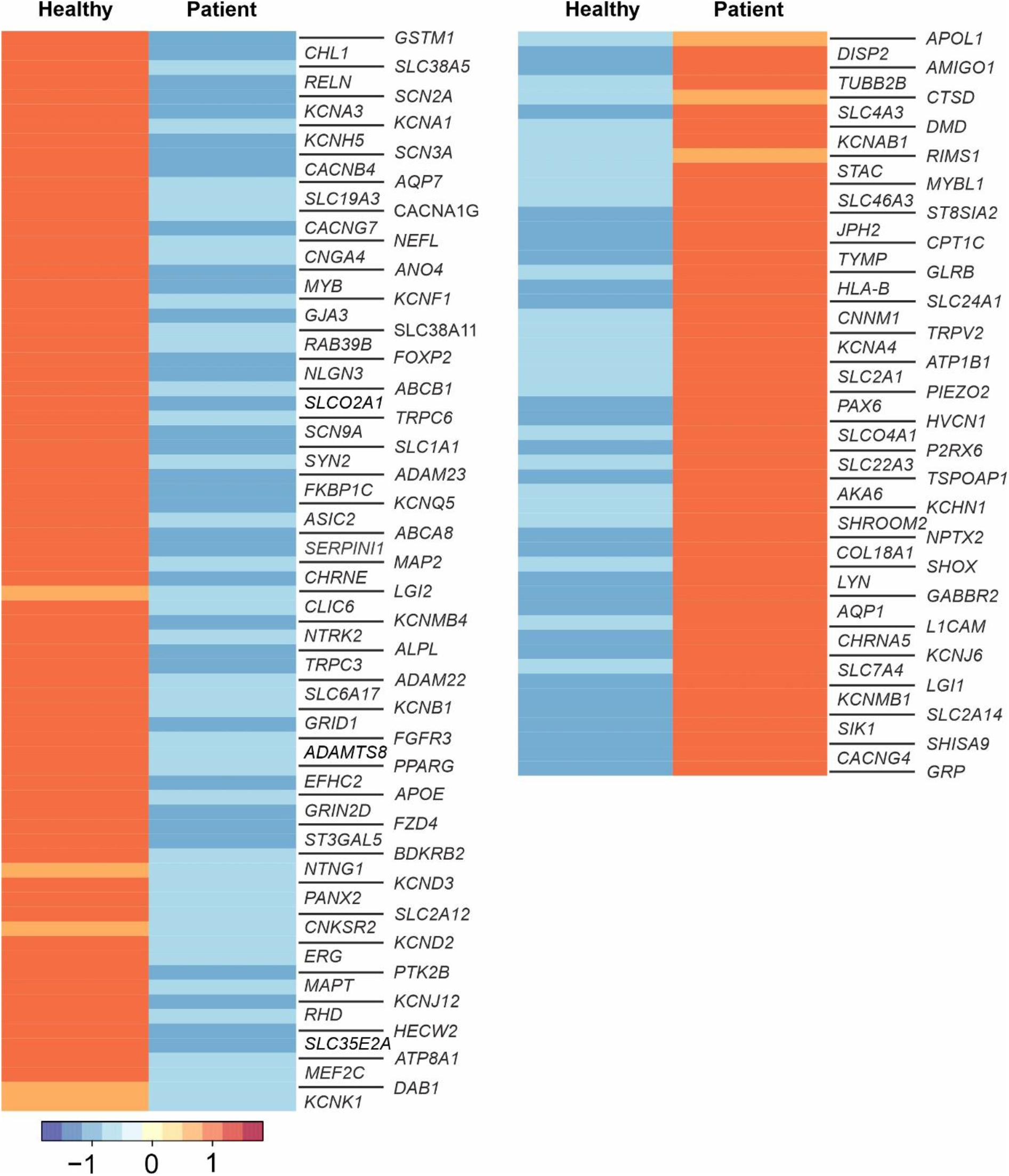
Differential expressed genes (DEGs) in patient-derived fibroblasts compared to fibroblasts from a healthy individuum. The average expression profiles from 3 replicates of healthy individuum and patient fibroblasts. Blue color shows downregulated genes and red color shows upregulated genes in patient-derived fibroblasts compared to healthy individuals.

**Figure S4.**
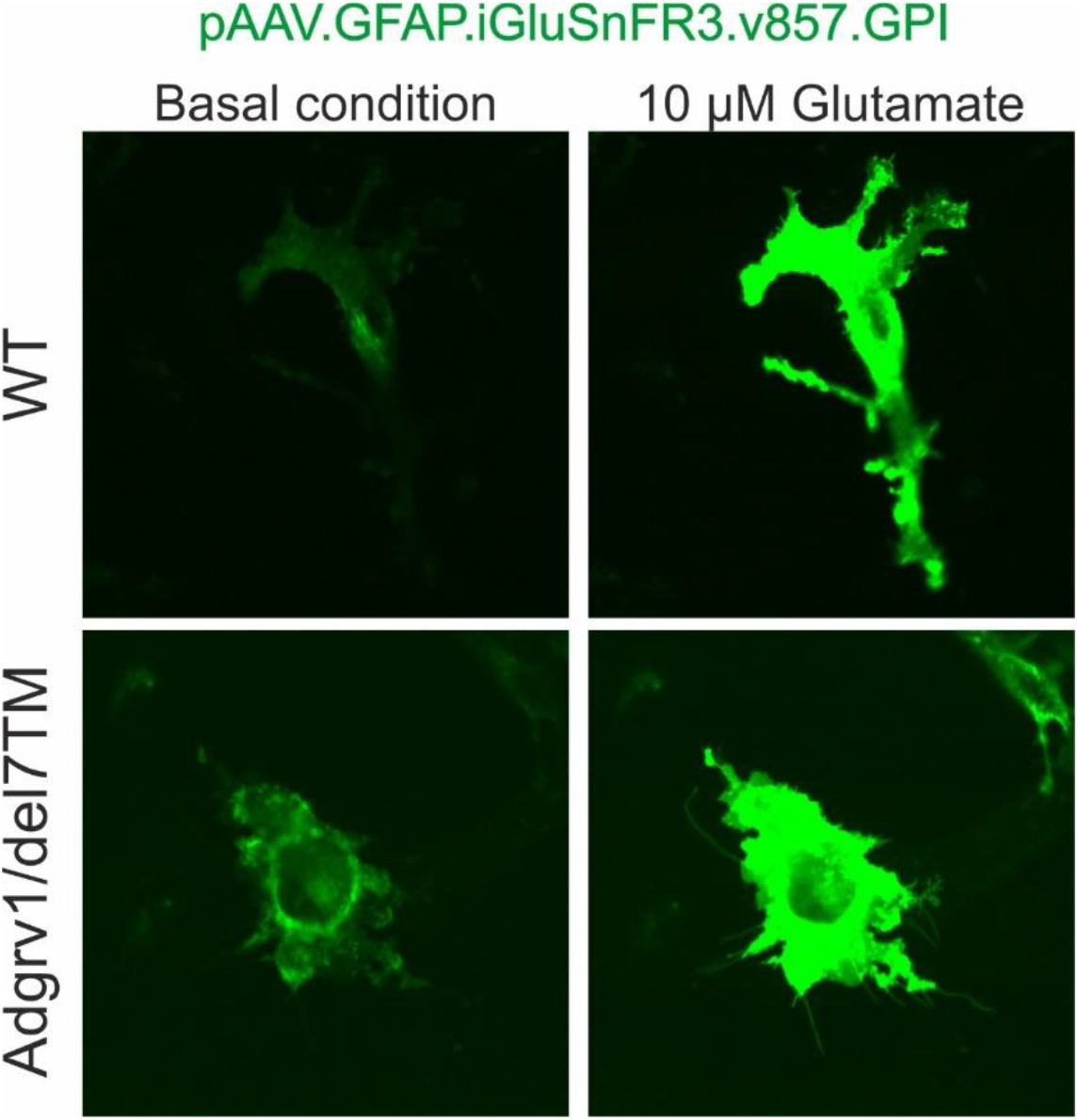
Images from live-cell imaging of pAAV.GFAP.iGluSnFR3.v857.GPI (green) expressing WT and Adgrv1/del7TM astrocytes. Time-lapse image sequences of pAAV.GFAP.iGluSnFR3.v857.GPI was recorded with 700 ms intervals for a total of 300 seconds. 10 µM glutamate was applied to the cells in 98th image of the sequence.

## References

Aggarwal A, Liu R, Chen Y, Ralowicz AJ, Bergerson SJ, Tomaska F, Mohar B, Hanson TL, Hasseman JP, Reep D, Tsegaye G, Yao P, Ji X, Kloos M, Walpita D, Patel R, Mohr MA, Tillberg PW, Looger LL, Marvin JS, Hoppa MB, Konnerth A, Kleinfeld D, Schreiter ER, Podgorski K. 2023. Glutamate indicators with improved activation kinetics and localization for imaging synaptic transmission. Nat Methods 20:925–934.

Alcoreza OB, Patel DC, Tewari BP, Sontheimer H. 2021. Dysregulation of Ambient Glutamate and Glutamate Receptors in Epilepsy: An Astrocytic Perspective. Front Neurol 12.

Allen NJ, Eroglu C. 2017. Cell Biology of Astrocyte-Synapse Interactions. Neuron 96:697–708.

Anderson GR, Maxeiner S, Sando R, Tsetsenis T, Malenka RC, Südhof TC. 2017. Postsynaptic adhesion GPCR latrophilin-2 mediates target recognition in entorhinal-hippocampal synapse assembly. J Cell Biol 216:3831–3846.

Arévalo AP, Lutz AK, Atanasova E, Boeckers TM. 2022. Trans-cardiac perfusion of neonatal mice and immunofluorescence of the whole body as a method to study nervous system development. PLoS One 17:1–8.

Arganda-Carreras I, Fernández-González R, Muñoz-Barrutia A, Ortiz-De-Solorzano C. 2010. 3D reconstruction of histological sections: Application to mammary gland tissue. Microsc Res Tech 73:1019–1029.

Armbruster M, Hanson E, Dulla CG. 2016. Glutamate clearance is locally modulated by presynaptic neuronal activity in the cerebral cortex. J Neurosci 36:10404–10415.

Bahi-Buisson N, El Sabbagh S, Soufflet C, Escande F, Boddaert N, Valayannopoulos V, Bellané-Chantelot C, Lascelles K, Dulac O, Plouin P, de Lonlay P. 2008. Myoclonic absence epilepsy with photosensitivity and a gain of function mutation in glutamate dehydrogenase. Seizure 17:658–664.

Baldwin KT, Eroglu C. 2017. Molecular mechanisms of astrocyte-induced synaptogenesis. Curr Opin Neurobiol 45:113–120.

Batiuk MY, Martirosyan A, Wahis J, de Vin F, Marneffe C, Kusserow C, Koeppen J, Viana JF, Oliveira JF, Voet T, Ponting CP, Belgard TG, Holt MG. 2020. Identification of region-specific astrocyte subtypes at single cell resolution. Nat Commun 11:1220.

Bindea G, Mlecnik B, Hackl H, Charoentong P, Tosolini M, Kirilovsky A, Fridman WH, Pagès F, Trajanoski Z, Galon J. 2009. ClueGO: A Cytoscape plug-in to decipher functionally grouped gene ontology and pathway annotation networks. Bioinformatics 25:1091–1093.

Boivin V, Reulet G, Boisvert O, Couture S, Elela SA, Scott MS. 2020. Reducing the structure bias of RNA-Seq reveals a large number of non-annotated non-coding RNA. Nucleic Acids Res 48:2271–2286.

Bolger AM, Lohse M, Usadel B. 2014. Trimmomatic: A flexible trimmer for Illumina sequence data. Bioinformatics 30:2114–2120.

Bonansco C, Fuenzalida M. 2016. Plasticity of hippocampal excitatory-inhibitory balance: Missing the synaptic control in the epileptic brain. Neural Plast 2016.

Bou Assi E, Nguyen DK, Rihana S, Sawan M. 2017. Towards accurate prediction of epileptic seizures: A review. Biomed Signal Process Control 34:144–157.

Broadhead MJ, Bonthron C, Waddington J, Smith W V., Lopez MF, Burley S, Valli J, Zhu F, Komiyama NH, Smith C, Grant SGN, Miles GB. 2022. Selective vulnerability of tripartite synapses in amyotrophic lateral sclerosis. Acta Neuropathol 143:471–486.

Chan F, Lax NZ, Voss CM, Aldana BI, Whyte S, Jenkins A, Nicholson C, Nichols S, Tilley E, Powell Z, Waagepetersen HS, Davies CH, Turnbull DM, Cunningham MO. 2019. The role of astrocytes in seizure generation: Insights from a novel in vitro seizure model based on mitochondrial dysfunction. Brain 142:391–411.

Chong D, Jones NC, Schittenhelm RB, Anderson A, Casillas-Espinosa PM. 2023. Multi-omics integration and epilepsy: Towards a better understanding of biological mechanisms. Prog Neurobiol 227:102480.

Chung W, Allen NJ, Eroglu C. 2015. Astrocytes Control Synapse Formation, Function, and Elimination. Cold Spring Harb Perspect Biol 7:a020370.

Cuellar-Santoyo AO, Ruiz-Rodríguez VM, Mares-Barbosa TB, Patrón-Soberano A, Howe AG, Portales-Pérez DP, Miquelajáuregui Graf A, Estrada-Sánchez AM. 2023. Revealing the contribution of astrocytes to glutamatergic neuronal transmission. Front Cell Neurosci 16:1– 15.

Raizen DM, Brooks-Kayal A, Steinkrauss L, Tennekoon GI, Stanley CA, Kelly A 2005. Central nervous system hyperexcitability associated with glutamate dehydrogenase gain of function mutations. J Pediatr 146:388–394.

Dzyubenko E, Rozenberg A, Hermann DM, Faissner A. 2016. Colocalization of synapse marker proteins evaluated by STED-microscopy reveals patterns of neuronal synapse distribution in vitro. J Neurosci Methods 273:149–159.

Falk MJ, Li D, Gai X, McCormick E, Place E, Lasorsa FM, Otieno FG, Hou C, Kim CE, Abdel-Magid N, Vazquez L, Mentch FD, Chiavacci R, Liang J, Liu X, Jiang H, Giannuzzi G, Marsh ED, Yiran G, Tian L, Palmieri F, Hakonarson H. 2014. AGC1 Deficiency Causes Infantile Epilepsy, Abnormal Myelination, and Reduced N-Acetylaspartate. In: JIMD Reports. Vol. 4. . p 77–85.

Ferreira TA, Blackman A V., Oyrer J, Jayabal S, Chung AJ, Watt AJ, Sjöström PJ, Van Meyel DJ. 2014. Neuronal morphometry directly from bitmap images. Nat Methods 11:982–984.

Gage GJ, Kipke DR, Shain W. 2012. Whole animal perfusion fixation for rodents. J Vis Exp:1– 9.

González-Reyes RE, Nava-Mesa MO, Vargas-Sánchez K, Ariza-Salamanca D, Mora-Muñoz L. 2017. Involvement of astrocytes in Alzheimer’s disease from a neuroinflammatory and oxidative stress perspective. Front Mol Neurosci 10:1–20.

Grüning B, Chilton J, Köster J, Dale R, Soranzo N, van den Beek M, Goecks J, Backofen R, Nekrutenko A, Taylor J. 2018. Practical Computational Reproducibility in the Life Sciences. Cell Syst 6:631–635.

Güler BE, Krzysko J, Wolfrum U. 2021. Isolation and culturing of primary mouse astrocytes for the analysis of focal adhesion dynamics. STAR Protoc 2.

Güler BE, Linnert J, Wolfrum U. 2023a. Monitoring paxillin in astrocytes reveals the significance of the adhesion GPCR VLGR1/ADGRV1 for focal adhesion assembly. Basic Clin Pharmacol Toxicol:1–12.

Güler BE, Linnert J, Wolfrum U. 2023b. Monitoring paxillin in astrocytes reveals the significance of the adhesion GPCR VLGR1/ADGRV1 for focal adhesion assembly. Basic Clin Pharmacol Toxicol:1–12.

Gürth CM, Dankovich TM, Rizzoli SO, D’Este E. 2020. Synaptic activity and strength are reflected by changes in the post-synaptic secretory pathway. Sci Rep 10:1–13.

Hamann J, Aust G, Araç D, Engel FB, Formstone C, Fredriksson R, Hall RA, Harty BL, Kirchhoff C, Knapp B, Krishnan A, Liebscher I, Lin HH, Martinelli DC, Monk KR, Peeters MC, Piao X, Prömel S, Schöneberg T, Schwartz TW, Singer K, Stacey M, Ushkaryov YA, Vallon M, Wolfrum U, Wright MW, Xu L, Langenhan T, Schiöth HB. 2015. International union of basic and clinical pharmacology. XCIV. adhesion G protein-coupled receptors. Pharmacol Rev 67:338–367.

Hayashi MK, Sato K, Sekino Y. 2022. Neurons Induce Tiled Astrocytes with Branches That Avoid Each Other. Int J Mol Sci 23.

Higashi K, Fujita A, Inanobe A, Tanemoto M, Doi K, Kubo T, Kurachi Y. 2001. An inwardly rectifying K+ channel, Kir4.1, expressed in astrocytes surrounds synapses and blood vessels in brain. Am J Physiol Cell Physiol 281:922–931.

Hillen AEJ, Heine VM. 2020. Glutamate Carrier Involvement in Mitochondrial Dysfunctioning in the Brain White Matter. Front Mol Biosci 7:1–8.

Holler S, Köstinger G, Martin KAC, Schuhknecht GFP, Stratford KJ. 2021. Structure and function of a neocortical synapse. Nature 591:111–116.

Ji T, Downs AW, Dorris L, Zhong N. 2023. De novo ADGRV1 variant in a patient with ictal asystole provides novel clues for increased risk of SUDEP. Acta Epileptologica 5.

Johannesen KM, Tümer Z, Weckhuysen S, Barakat TS, Bayat A. 2023. Solving the unsolved genetic epilepsies: Current and future perspectives. Epilepsia 64:3143–3154.

Kim D, Paggi JM, Park C, Bennett C, Salzberg SL. 2019. Graph-based genome alignment and genotyping with HISAT2 and HISAT-genotype. Nat Biotechnol 37:907–915.

Kinoshita T, Fujita M. 2016. Biosynthesis of GPI-anchored proteins: Special emphasis on GPI lipid remodeling. J Lipid Res 57:6–24.

Knapp B, Roedig J, Boldt K, Krzysko J, Horn N, Ueffing M, Wolfrum U. 2019. Affinity proteomics identifies novel functional modules related to adhesion GPCRs. Ann N Y Acad Sci 1456:144–167.

Knapp B, Roedig J, Roedig H, Krzysko J, Horn N, Güler BE, Kusuluri DK, Yildirim A, Boldt K, Ueffing M, Liebscher I, Wolfrum U. 2022. Affinity Proteomics Identifies Interaction Partners and Defines Novel Insights into the Function of the Adhesion GPCR VLGR1/ADGRV1. Molecules 27.

Kritis AA, Stamoula EG, Paniskaki KA, Vavilis TD. 2015. Researching glutamate – induced cytotoxicity in different cell lines: A comparative/collective analysis/study. Front Cell Neurosci 9:1–18.

Krzysko J, Maciag F, Mertens A, Güler BE, Linnert J, Boldt K, Ueffing M, Nagel-Wolfrum K, Heine M, Wolfrum U. 2022. The Adhesion GPCR VLGR1/ADGRV1 Regulates the Ca2+ Homeostasis at Mitochondria-Associated ER Membranes. Cells 11:1–20.

Kusuluri DK, Güler BE, Knapp B, Horn N, Boldt K, Ueffing M, Aust G, Wolfrum U. 2021. Adhesion G protein-coupled receptor VLGR1/ADGRV1 regulates cell spreading and migration by mechanosensing at focal adhesions. iScience 24.

Langenhan T. 2020. Adhesion G protein–coupled receptors—Candidate metabotropic mechanosensors and novel drug targets. Basic Clin Pharmacol Toxicol 126:5–16.

Langer J, Gerkau NJ, Derouiche A, Kleinhans C, Moshrefi-Ravasdjani B, Fredrich M, Kafitz KW, Seifert G, Steinhäuser C, Rose CR. 2017. Rapid sodium signaling couples glutamate uptake to breakdown of ATP in perivascular astrocyte endfeet. Glia 65:293–308.

Lechan RM, Fekete C. 2006. Chapter 12: The TRH neuron: a hypothalamic integrator of energy metabolism. Prog Brain Res 153:209–235.

Lefèvre G, Michel V, Weil D, Lepelletier L, Bizard E, Wolfrum U, Hardelin JP, Petit C. 2008. A core cochlear phenotype in USH1 mouse mutants implicates fibrous links of the hair bundle in its cohesion, orientation and differential growth. Development 135:1427–1437.

Leng X, Zhang T, Guan Y, Tang M. 2022. Genotype and phenotype analysis of epilepsy caused by ADGRV1 mutations in Chinese children. Seizure 103:108–114.

Li H, Handsaker B, Wysoker A, Fennell T, Ruan J, Homer N, Marth G, Abecasis G, Durbin R. 2009. The Sequence Alignment/Map format and SAMtools. Bioinformatics 25:2078–2079.

Liao Y, Smyth GK, Shi W. 2014. FeatureCounts: An efficient general purpose program for assigning sequence reads to genomic features. Bioinformatics 30:923–930.

Liu JYW, Dzurova N, Al-Kaaby B, Mills K, Sisodiya SM, Thom M. 2020. Granule Cell Dispersion in Human Temporal Lobe Epilepsy: Proteomics Investigation of Neurodevelopmental Migratory Pathways. Front Cell Neurosci 14.

Love MI, Huber W, Anders S. 2014. Moderated estimation of fold change and dispersion for RNA-seq data with DESeq2. Genome Biol 15:1–21.

Maerker T, van Wijk E, Overlack N, Kersten FFJ, Mcgee J, Goldmann T, Sehn E, Roepman R, Walsh EJ, Kremer H, Wolfrum U. 2008. A novel Usher protein network at the periciliary reloading point between molecular transport machineries in vertebrate photoreceptor cells. Hum Mol Genet 17:71–86.

Mahmoud S, Gharagozloo M, Simard C, Amrani A, Gris D. 2019. NLRX1 enhances glutamate uptake and inhibits glutamate release by astrocytes. Cells 8:1–16.

Mata G, Heras J, Morales M, Romero A, Rubio J. 2016. SynapCountJ: A tool for analyzing synaptic densities in neurons. BIOIMAGING 2016 - 3rd International Conference on Bioimaging, Proceedings; Part of 9th International Joint Conference on Biomedical Engineering Systems and Technologies, BIOSTEC 2016:25–31.

McMillan DR, White PC. 2004. Loss of the transmembrane and cytoplasmic domains of the very large G-protein-coupled receptor-1 (VLGR1 or Mass1) causes audiogenic seizures in mice. Mol Cell Neurosci 26:322–329.

McMillan DR, White PC. 2010. Studies on the very large G protein-coupled receptor: from initial discovery to determining its role in sensorineural deafness in higher animals. Adv Exp Med Biol 706:76–86.

Michalski N, Michel V, Bahloul A, Lefèvre G, Barral J, Yagi H, Chardenoux S, Weil D, Martin P, Hardelin JP, Sato M, Petit C. 2007. Molecular characterization of the ankle-link complex in cochlear hair cells and its role in the hair bundle functioning. Journal of Neuroscience 27:6478– 6488.

Michinaga S, Koyama Y. 2019. Dual roles of astrocyte-derived factors in regulation of blood-brain barrier function after brain damage. Int J Mol Sci 20:1–22.

Mölder F, Jablonski KP, Letcher B, Hall MB, Tomkins-Tinch CH, Sochat V, Forster J, Lee S, Twardziok SO, Kanitz A, Wilm A, Holtgrewe M, Rahmann S, Nahnsen S, Köster J. 2021. Sustainable data analysis with Snakemake [version 2; peer review: 2 approved]. F1000Res 10:1–29.

Myers KA, Nasioulas S, Boys A, McMahon JM, Slater H, Lockhart P, Sart D du, Scheffer IE. 2018a. ADGRV1 is implicated in myoclonic epilepsy. Epilepsia 59:381–388.

Nissen JD, Pajęcka K, Stridh MH, Skytt DM, Waagepetersen HS. 2015. Dysfunctional TCA-Cycle Metabolism in Glutamate Dehydrogenase Deficient Astrocytes. Glia 63:2313–2326.

Onaolapo AY, Onaolapo OJ. 2020. Peripheral and Central Glutamate Dyshomeostasis in Neurodegenerative Disorders. Curr Neuropharmacol 19:1069–1089.

Ouyang YB, Voloboueva LA, Xu LJ, Giffard RG. 2007. Selective dysfunction of hippocampal CA1 astrocytes contributes to delayed neuronal damage after transient forebrain ischemia. Journal of Neuroscience 27:4253–4260.

Papageorgiou IE, Valous NA, Lahrmann B, Janova H, Klaft Z, Koch A, Schneider UC, Vajkoczy P, Heppner FL, Grabe N, Halama N, Heinemann U, Kann O. 2018. Astrocytic glutamine synthetase is expressed in the neuronal somatic layers and down-regulated proportionally to neuronal loss in the human epileptic hippocampus. Glia 66:920–933.

Perea G, Navarrete M, Araque A. 2009. Tripartite synapses: astrocytes process and control synaptic information. Trends Neurosci 32:421–431.

Perez-Catalan NA, Doe CQ, Ackerman SD. 2021. The role of astrocyte-mediated plasticity in neural circuit development and function. Neural Dev 16:1–14.

Piepgras J, Rohrbeck A, Just I, Bittner S, Ahnert-Hilger G, Höltje M. 2022. Enhancement of Phosphorylation and Transport Activity of the Neuronal Glutamate Transporter Excitatory Amino Acid Transporter 3 by C3bot and a 26mer C3bot Peptide. Front Cell Neurosci 16:1–14.

Poduri A, Heinzen EL, Chitsazzadeh V, Lasorsa FM, Elhosary PC, LaCoursiere CM, Martin E, Yuskaitis CJ, Hill RS, Atabay KD, Barry B, Partlow JN, Bashiri FA, Zeidan RM, Elmalik SA, Kabiraj MMU, Kothare S, Stödberg T, McTague A, Kurian MA, Scheffer IE, Barkovich AJ, Palmieri F, Salih MA, Walsh CA. 2013. SLC25A22 is a novel gene for migrating partial seizures in infancy. Ann Neurol 74:873–882.

Robinson MB, Lee ML, DaSilva S. 2020. Glutamate Transporters and Mitochondria: Signaling, Co-compartmentalization, Functional Coupling, and Future Directions. Neurochem Res 45:526–540.

Rose CR, Felix L, Zeug A, Dietrich D, Reiner A, Henneberger C. 2018. Astroglial glutamate signaling and uptake in the hippocampus. Front Mol Neurosci 10:1–20.

Rose J, Brian C, Pappa A, Panayiotidis MI, Franco R. 2020. Mitochondrial Metabolism in Astrocytes Regulates Brain Bioenergetics, Neurotransmission and Redox Balance. Front Neurosci 14:1–20.

Rouillard AD, Gundersen GW, Fernandez NF, Wang Z, Monteiro CD, McDermott MG, Ma’ayan A. 2016. The harmonizome: a collection of processed datasets gathered to serve and mine knowledge about genes and proteins. Database (Oxford) 2016:1–16.

Sachs N, Wechsberg O, Landau YE, Krause I, Elgali II, Darawshe M, Shomron N, Lidzbarsky G, Orenstein N. 2023. A novel SLC25A13 gene splice site variant causes Citrin deficiency in an infant. Gene 874:147483.

Saleh M, Helmi M, Yacop B. 2020. A Novel Nonsense Gene Variant Responsible for Early Infantile Epileptic Encephalopathy Type 39: Case Report. Pak J Biol Sci 23:973–976.

Sidoryk-Wegrzynowicz M, Wegrzynowicz M, Lee E, Bowman AB, Aschner M. 2011. Role of Astrocytes in Brain Function and Disease. Toxicol Pathol 39:115–123.

de Sousa DMB, Benedetti A, Altendorfer B, Mrowetz H, Unger MS, Schallmoser K, Aigner L, Kniewallner KM. 2023. Immune-mediated platelet depletion augments Alzheimer’s disease neuropathological hallmarks in APP-PS1 mice. Aging 15:630–649.

Steinhäuser C, Seifert G, Bedner P. 2012. Astrocyte dysfunction in temporal lobe epilepsy: K + channels and gap junction coupling. Glia 60:1192–1202.

Szklarczyk D, Franceschini A, Wyder S, Forslund K, Heller D, Huerta-Cepas J, Simonovic M, Roth A, Santos A, Tsafou KP, Kuhn M, Bork P, Jensen LJ, Von Mering C. 2015. STRING v10: Protein-protein interaction networks, integrated over the tree of life. Nucleic Acids Res 43:D447–D452.

Trabelsi Y, Amri M, Becq H, Molinari F, Aniksztejn L. 2017. The conversion of glutamate by glutamine synthase in neocortical astrocytes from juvenile rat is important to limit glutamate spillover and peri/extrasynaptic activation of NMDA receptors. Glia 65:401–415.

Wang Y, Fan X, Zhang W, Zhang C, Wang J, Jiang T, Wang L. 2015. Deficiency of very large G-protein-coupled receptor-1 is a risk factor of tumor-related epilepsy: a whole transcriptome sequencing analysis. J Neurooncol 121:609–616.

Weston MD, Luijendijk MWJ, Humphrey KD, Möller C, Kimberling WJ. 2004. Mutations in the VLGR1 Gene Implicate G-Protein Signaling in the Pathogenesis of Usher Syndrome Type II. Am J Hum Genet 74:357–366.

Wingett SW, Andrews S. 2018. Fastq screen: A tool for multi-genome mapping and quality control. F1000Res 7:1–14.

Wolfrum U. 1991. Distribution of F-actin in the compound eye of the blowfly, Calliphora erythrocephala (Diptera, Insecta). Cell Tissue Res 263:399–403.

Wu Z, Deshpande T, Henning L, Bedner P, Seifert G, Steinhäuser C. 2021. Cell death of hippocampal CA1 astrocytes during early epileptogenesis. Epilepsia 62:1569–1583.

Young K, Morrison H. 2018. Quantifying microglia morphology from photomicrographs of immunohistochemistry prepared tissue using imagej. JOVE 2018:1–9.

Zhang Y, Sloan SA, Clarke LE, Caneda C, Plaza CA, Blumenthal PD, Vogel H, Steinberg GK, Edwards MSB, Li G, Duncan JA, Cheshier SH, Shuer LM, Chang EF, Grant GA, Gephart MGH, Barres BA. 2016. Purification and Characterization of Progenitor and Mature Human Astrocytes Reveals Transcriptional and Functional Differences with Mouse. Neuron 89:37–53.

Zhou P, Meng H, Liang X, Lei X, Zhang J, Bian W, He N, Lin Z, Song X, Zhu W, Hu B, Li B, Yan L, Tang B, Su T, Liu H, Mao Y, Zhai Q, Yi Y. 2022. ADGRV1 Variants in Febrile Seizures/Epilepsy With Antecedent Febrile Seizures and Their Associations With Audio-Visual Abnormalities. Front Mol Neurosci 15.

Zhou Y, Wang X, Tzingounis A V., Danbolt NC, Larsson HP. 2014. EAAT2 (GLT-1; slc1a2) glutamate transporters reconstituted in liposomes argues against heteroexchange being substantially faster than net uptake. Journal of Neuroscience 34:13472–13485.

