## Supplementary Figures for "The adhesion GPCR ADGRV1 controls glutamate homeostasis in hippocampal astrocytes supporting neuron development: first insights into to pathophysiology of *ADGRV1*-associated epilepsy"

**Supplemental Figures:**

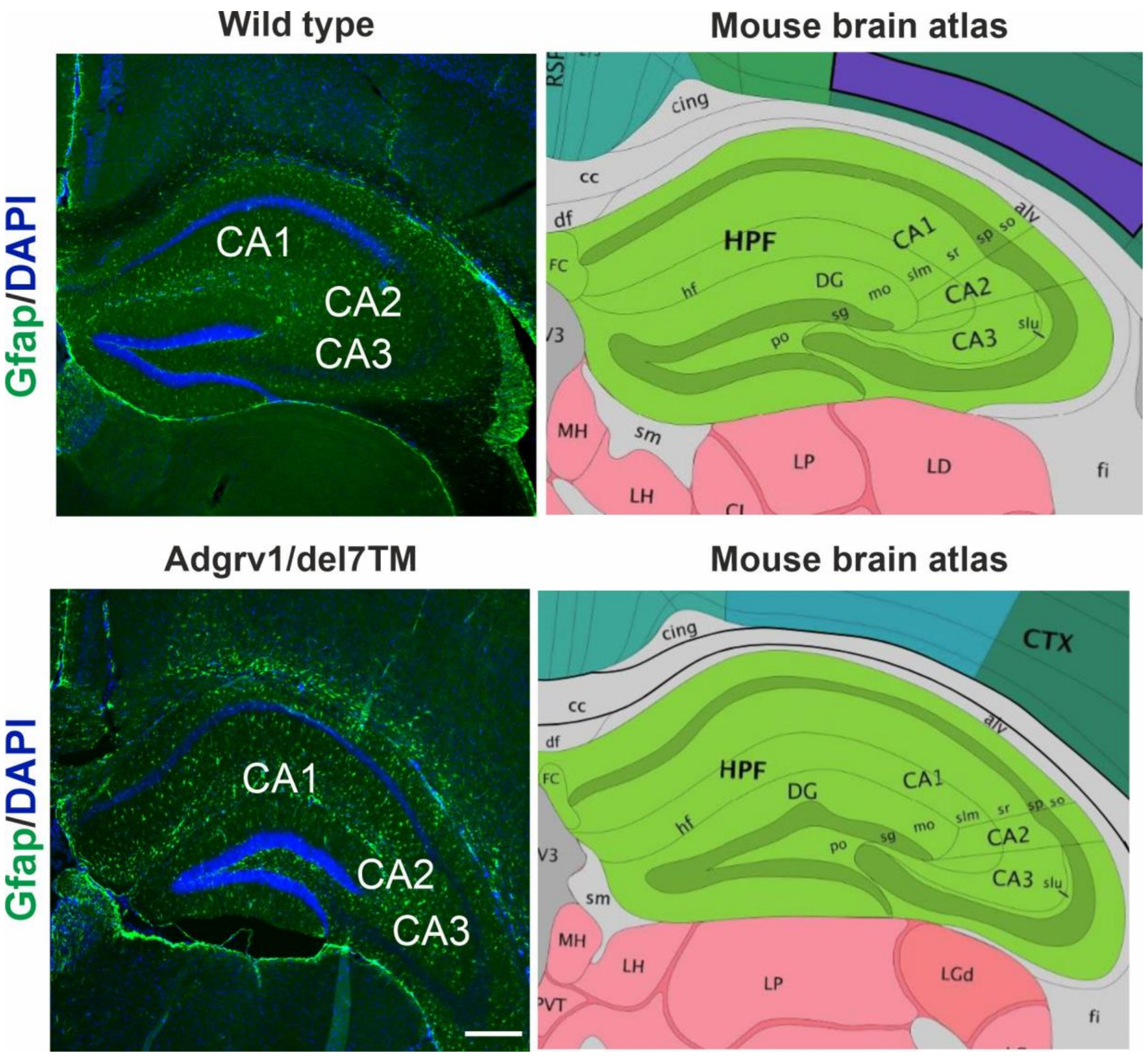


**Figure S1. Reference coronal brain sections from mouse brain atlas for the identification of hippocampus subregions.**

The subregions of the mouse hippocampus sections were identified using the Allen mouse brain atlas (https://mouse.brain-map.org/) during the image analysis. Scale bar: 25 µm.

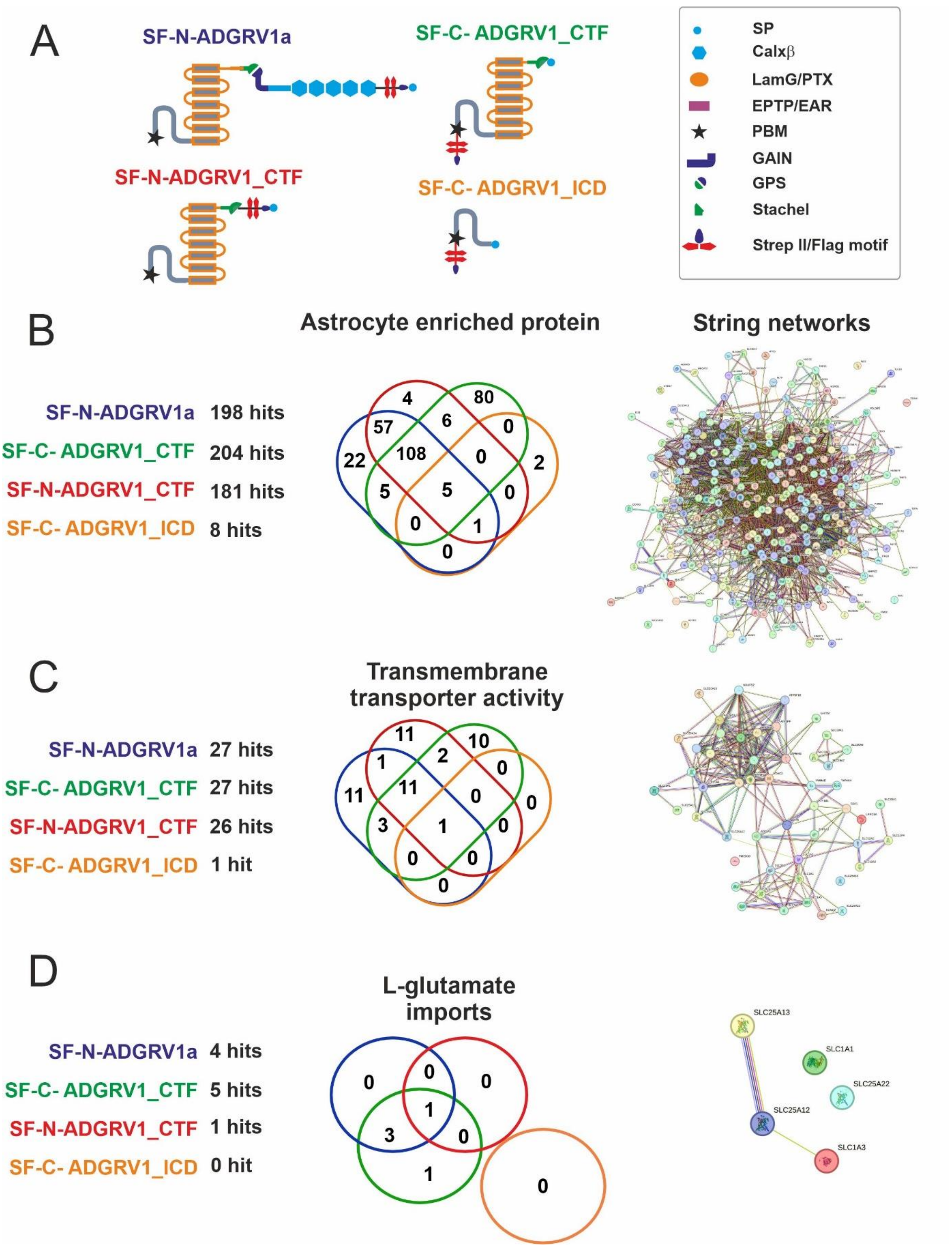
A
SF-N-ADGRV1a
SF-C- ADGRV1_CTF
SF-N-ADGRV1_CTF
SF-C- ADGRV1_ICD
SP
Calxβ
LamG/PTX
EPTP/EAR
PBM
GAIN
GPS
Stachel
Strep II/Flag motif
B
Astrocyte enriched protein
String networks
SF-N-ADGRV1a 198 hits
SF-C- ADGRV1_CTF 204 hits
SF-N-ADGRV1_CTF 181 hits
SF-C- ADGRV1_ICD 8 hits
4
80
57
6
0
22
108
0
2
5
5
0
0
1
0
C
Transmembrane transporter activity
SF-N-ADGRV1a 27 hits
SF-C- ADGRV1_CTF 27 hits
SF-N-ADGRV1_CTF 26 hits
SF-C- ADGRV1_ICD 1 hit
11
10
1
2
0
11
11
0
0
3
1
0
0
0
0
D
L-glutamate imports
SF-N-ADGRV1a 4 hits
SF-C- ADGRV1_CTF 5 hits
SF-N-ADGRV1_CTF 1 hits
SF-C- ADGRV1_ICD 0 hit
0
0
0
1
3
0
1
0
SLC25A13
SLC1A1
SLC25A22
SLC25A12
SLC1A3

**Figure S2. TAP analysis reveals a complex protein network related to ADGRV1 protein**

**(A)** Different Strep II/FLAG (SF)-tagged ADGRV1 constructs used as a prey in Tanden affinity purification (TAP) to revealing potential interaction partners of ADGRV1 using HEK293T cells. ADGRV1 constructs were tagged from C or N terminals to eliminate false binding partners and changes in receptor structure. (**B**) GO term analysis revealed astrocyte enriched protein in ADGRV1 TAP analysis. The Venn-diagram shows a high overlap in astrocyte enriched proteins in 4 different ADGRV1 constructs, and string network analysis shows interactions. (**C**) Transmembrane transporter activity related proteins highly overlapped in SF-N-ADGRV1a, SF-N-ADGRV1_CTF and SF-C-ADGRV1 preys. (**D**) L-glutamate import related proteins have enriched in SF-N-ADGRV1a and SF-C-ADGRV1 preys.

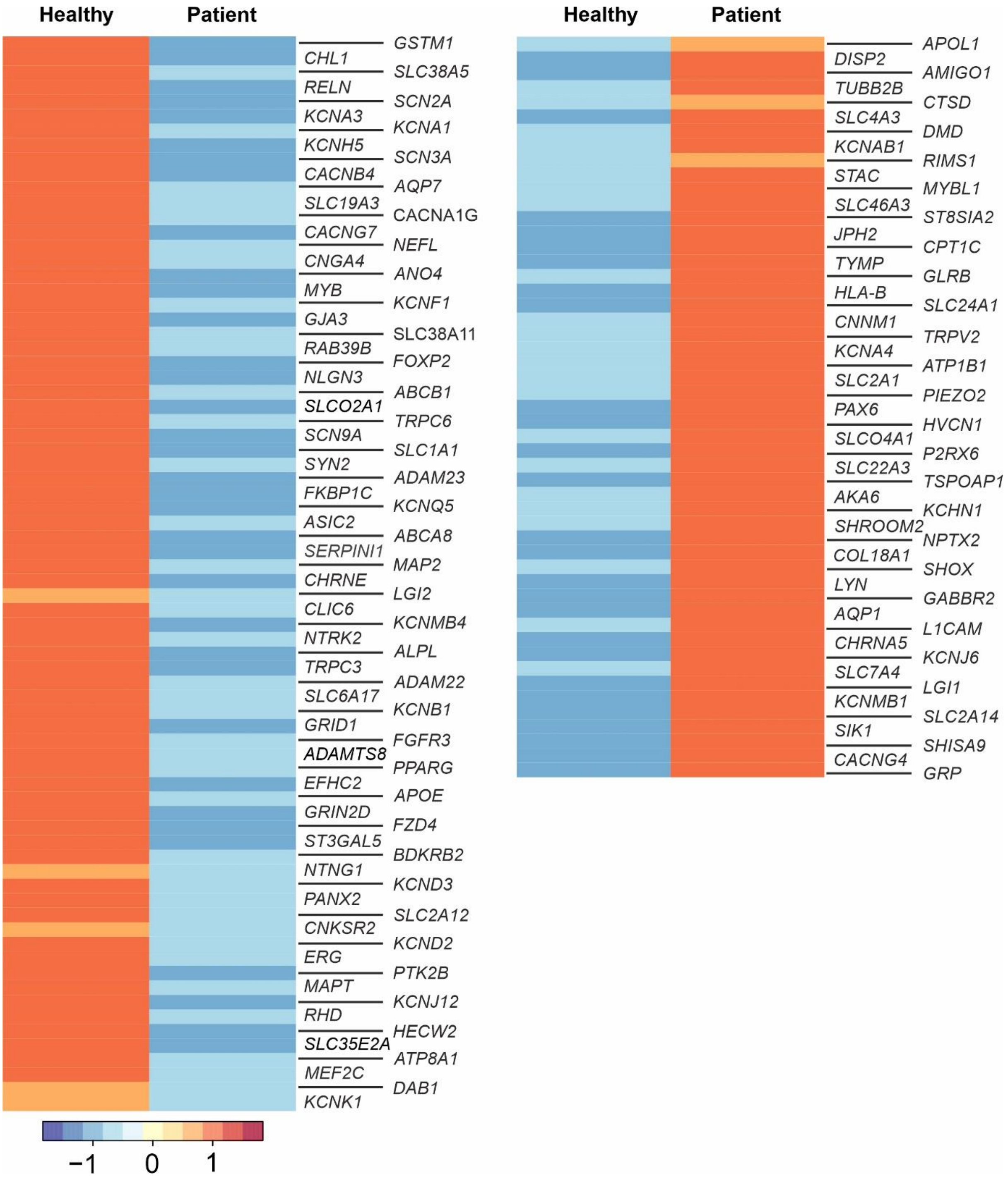


**Figure S3. Differential expressed genes (DEGs) in patient-derived fibroblasts compared to fibroblasts from a healthy individuum.** The average expression profiles from 3 replicates of healthy individuum and patient fibroblasts. Blue color shows downregulated genes and red color shows upregulated genes in patient-derived fibroblasts compared to healthy individuals.

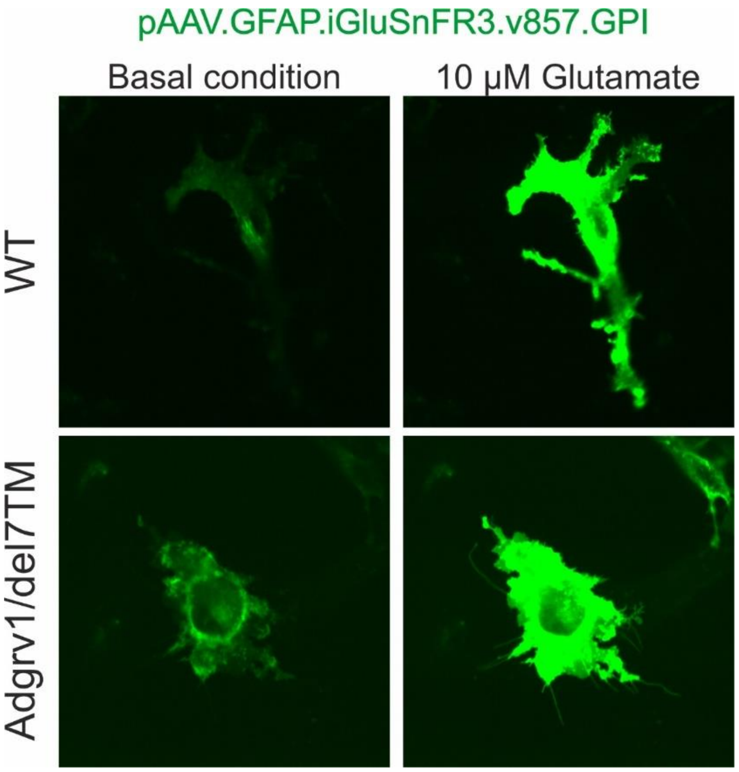


**Figure S4. Images from live-cell imaging of pAAV.GFAP.iGluSnFR3.v857.GPI (green) expressing WT and Adgrv1/del7TM astrocytes.**

Time-lapse image sequences of pAAV.GFAP.iGluSnFR3.v857.GPI was recorded with 700 ms intervals for a total of 300 seconds. 10 µM glutamate was applied to the cells in 98th image of the sequence.

Uncropped Western blots: Güler et al.

To Figure 2A

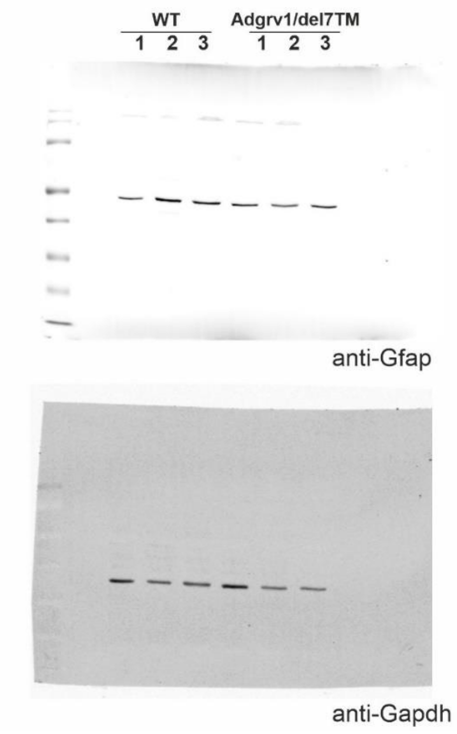


To Figure 2B

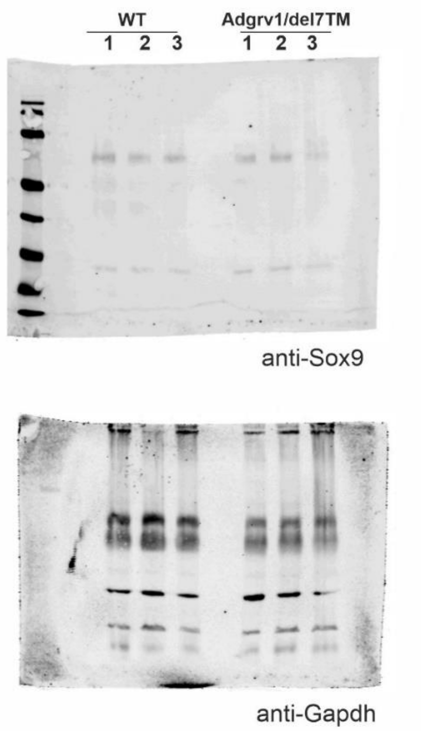

To Figure 6

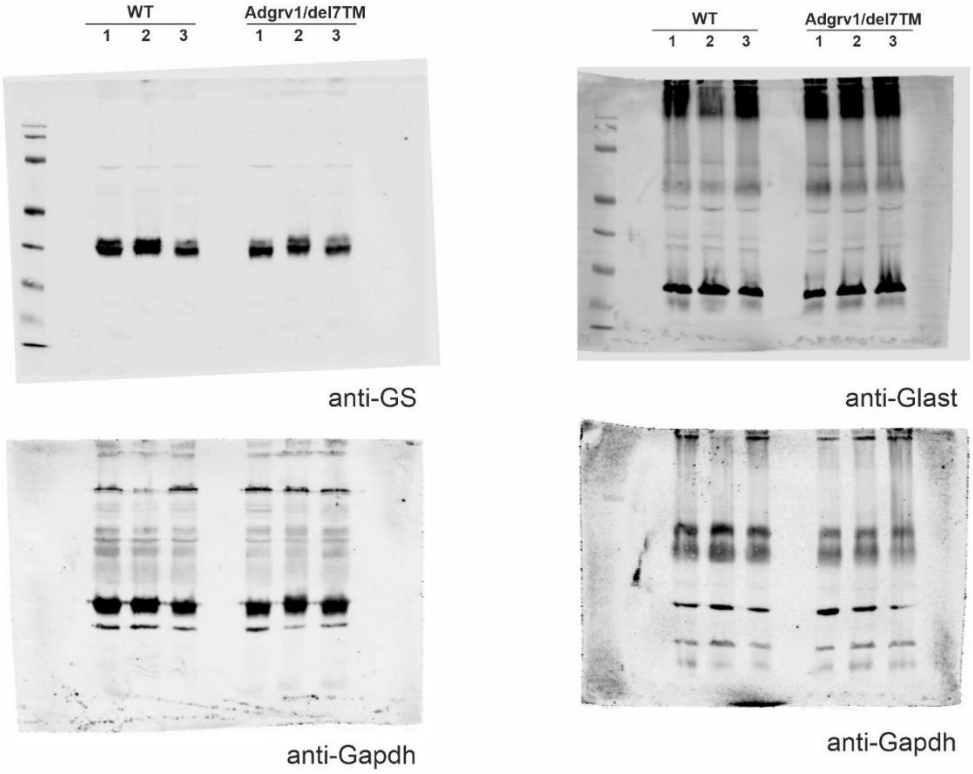
WT
Adgrv1/del7TM
1 2 3
1 2 3
anti-GS
anti-Glast
anti-Gapdh
anti-Gapdh
